# Synthetic data enables human-grade microtubule analysis with foundation models for segmentation

**DOI:** 10.64898/2026.01.09.698597

**Authors:** Mario Koddenbrock, Justus Westerhoff, Dominik Fachet, Simone Reber, Felix Gers, Erik Rodner

## Abstract

Studying microtubules (MTs) and their mechanical properties is central to understanding intracellular transport, cell division, and drug action. While important, experts still need to spend many hours manually segmenting these filamentous structures. The suitability of state-of-the-art methods for this task cannot be systematically assessed, as large-scale labeled datasets are missing. We address this gap by presenting the synthetic dataset SynthMT, produced by tuning a novel image generation pipeline on real-world interference reflection microscopy (IRM) frames of *in vitro* reconstituted MTs without requiring human annotations. In our benchmark, we evaluate nine fully automated methods for MT analysis in both zero- and Hyperparameter Optimization (HPO)-based few-shot settings. Across both settings, classical algorithms and current foundation models still struggle to achieve the accuracy required for biological downstream analysis on *in vitro* MT IRM images that humans perceive as visually simple. However, a notable exception is the recently introduced SAM3 model. After HPO on only ten random SynthMT images, its text-prompted version SAM3Text achieves near-perfect and in some cases super-human performance on unseen, real data. This indicates that fully automated MT segmentation has become feasible when method configuration is effectively guided by synthetic data. To enable progress, we publicly release the dataset, the generation pipeline, and the evaluation framework at DATEXIS.github.io/SynthMT-project-page.

**Author summary:** Understanding the behavior of microtubules — stiff filaments inside cells — is essential for studying fundamental cell biological processes and for developing therapies for diseases such as cancer and neurodegenerative conditions. Yet, analyzing microtubule images is slow and labor-intensive, as researchers must manually trace these thin, overlapping filaments, which can take hours and is prone to errors. We therefore asked whether current fully automated segmentation methods are ready to replace manual microtubule analysis, and how synthetic data can be used to rigorously evaluate and improve them.

To address these questions, we created a synthetic dataset that mimics real microtubule images by capturing the appearance and variability of real microscopy. Importantly, this dataset can be generated without any manual annotations. Using this dataset, we evaluated a range of segmentation methods and found that most of them struggled to accurately identify filaments. However, we discovered that a recent foundation model, when guided by a simple text instruction and tuned on only a few synthetic images, can achieve near-perfect, human-level performance on previously unseen real microtubule imaging data.

Our work demonstrates that fully automated microtubule analysis is now possible and provides a reproducible framework that other researchers can use to evaluate and improve their methods. This opens the door to faster, more consistent studies of microtubules, ultimately accelerating discoveries in cell biology and therapeutic research.

## 1 Introduction

Microtubules (MTs) are cytoskeletal filaments essential for cellular processes such as chromosome segregation, intracellular transport, and cell motility. They are formed by head-to-tail polymerization of *αβ*-tubulin heterodimers into protofilaments that laterally associate to form a hollow cylinder with a diameter of about 25 *nm* [1]. Their plus ends exhibit *dynamic instability*, stochastically switching between growth and shrinkage, a behavior exquisitely sensitive to regulatory proteins and small molecules. This sensitivity makes MTs prime targets in drug discovery: widely used chemotherapeutics (e.g., taxanes [2], vinca alkaloids [3]) exploit destabilization, while emerging neuroprotective strategies aim to stabilize MT dynamics in neurodegenerative diseases [4]. To investigate how candidate compounds, cofactors, or post-translational modifications alter growth behavior, curvature, and length distributions, MTs are often reconstituted *in vitro* from stabilized nucleation seeds and visualized using microscopy techniques like total internal reflection fluorescence (TIRF) or interference reflection microscopy (IRM). A key step in such analyses is the accurate quantification of filament properties (count, length, curvature), which requires precise instance segmentation. However, manual annotation of MT images is labor-intensive and error-prone, creating a significant bottleneck in experimental and preclinical workflows.

Automated segmentation methods offer a promising solution, but their development and evaluation are hampered by a major challenge: the lack of large-scale, publicly available, and annotated datasets for *in vitro* MT images that resemble the typical noise of biological imaging data. This data scarcity has been repeatedly noted [5–7], yet no comprehensive benchmark exists.

While general-purpose segmentation models, including recent foundation models like CellSAM [10], *µ*SAM [11], and Cellpose-SAM [12], have shown impressive zero-shot performance on various microscopy datasets, they are primarily designed for round objects such as nuclei or whole cells. Their applicability to filamentous structures such as MTs remains largely unproven, and they often lack the accuracy necessary for reliable downstream biological analysis, especially when dealing with out-of-distribution data [13–16]. This raises a central question: **Are current fully automated methods ready to replace manual analysis for MT segmentation?** As sufficiently advanced methods often critically depend on their hyperparameters, we also ask to what extent synthetic data that mimics real microscopy images can enable few-shot learning through Hyperparameter Optimization (HPO), thereby improving generalization to real-world MT data.

This paper addresses these questions by introducing SynthMT, a synthetic benchmark designed to systematically evaluate the readiness of segmentation methods for automated MT analysis in zero- and few-shot settings (see Fig. 1).

**Figure 1.**
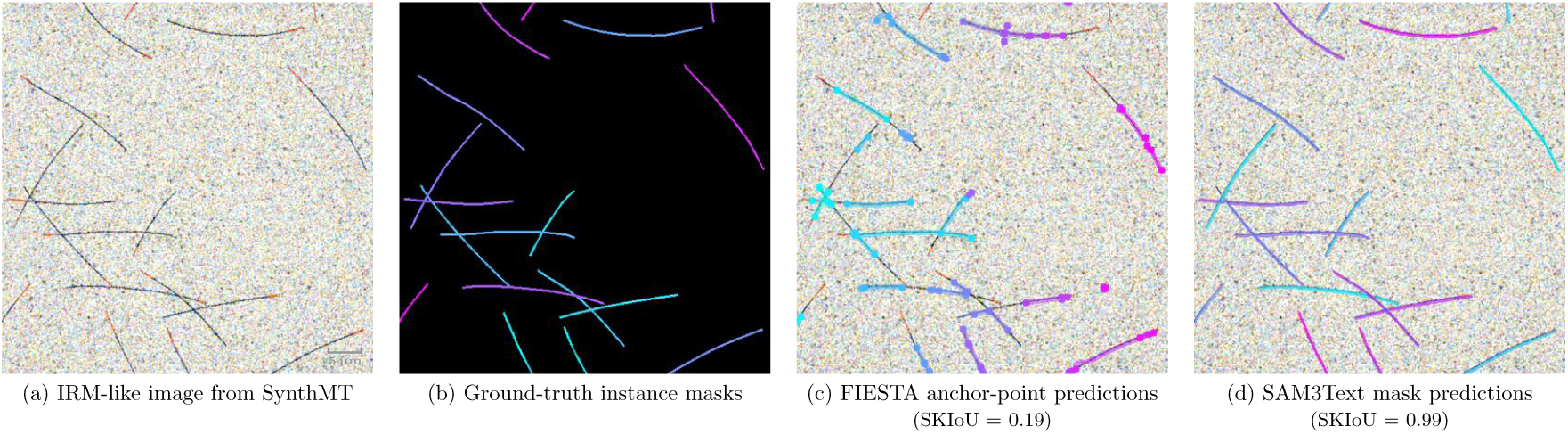
The SynthMT instance segmentation benchmark evaluates methods on synthetic interference reflection microscopy (IRM)-like images containing microtubules (MTs). (a) Synthetic image mimicking IRM of *in vitro* reconstituted MTs nucleated from fixed seeds (visualized in red), reproducing key mechanical and geometrical properties such as filament length and curvature. (b) Our pipeline generates accompanying ground-truth instance masks that enable quantitative evaluation. (c) The classical FIESTA [8] algorithm predicts anchor points for each instance (for visual clarity, only the first and last point of each instance are shown), which we connect through splines. The example demonstrates typical failure modes: filament fragmentation (single MTs split into multiple instances), incomplete segmentation, and artifacts at intersections. (d) SAM3 [9] guided by a simple text prompt (“*thin line*”) produces precise, human-grade segmentation, accurately tracing intersecting MTs. This is supported by its high Skeleton Intersection over Union (SKIoU) for this specific image.

Our main contributions and results are:

- **Synthetic data generation pipeline replicating real images.** Our publicly available pipeline generates realistic MT images along with corresponding ground-truth annotations. For tuning its internal parameters, it only requires real MT images (IRM or total internal reflection fluorescence (TIRF)) without the need for human annotations.
- **Synthetic benchmark dataset (**SynthMT). We apply this generation pipeline to real IRM images to construct and release the SynthMT dataset including instance masks for each MT. Its biological plausibility is assessed through a study with domain experts.
- **Comprehensive evaluation.** We conduct a reproducible benchmark evaluation of classical and state-of-the-art segmentation methods on SynthMT, including zero-shot and HPO-based few-shot adaptation experiments, establishing quantitative baselines.
- **Enabling fully automated segmentation.** We show that the newly released foundation model SAM3 achieves human-grade performance on MT images when parameter-optimized via SynthMT. Our synthetic data thus makes it possible to adapt this general-purpose model to the specific domain of *in vitro* MT microscopy, establishing a practical route toward fully automated segmentation for large-scale experiments and subsequent geometric analyses.

To facilitate progress, we publicly release the generation pipeline, dataset, and evaluation framework at DATEXIS.github.io/SynthMT-project-page, providing the community with tools to benchmark existing methods and accelerate method development.

## 2 Related work

### Synthetic datasets for microscopy

Synthetic data has recently been explored as a way to overcome the lack of annotated microscopy datasets. For example, AnyStar [17] employs a generative model to produce synthetic 3D training volumes of star-convex objects used to train StarDist [18] models, enabling zero-shot generalization across organisms without retraining. While powerful for spherical targets such as nuclei, the star-convex shape assumption makes this approach inapplicable to filamentous structures such as microtubules (MTs), where continuity and curvature priors are essential.

The concurrently released MicSim FluoMT [19] dataset represents an important step toward synthetic benchmarks for *in vivo* MTs. It provides microscopy-like images together with ground-truth segmentation masks, generated using Cytosim [20] for filament dynamics and ConfocalGN [21] for imaging rendering (see Fig. A.5a for examples). Their evaluation focuses on architectures trained entirely from scratch, but does not address the performance of existing foundation models.

Saguy et al. [22] propose a diffusion-based generative model for MT imaging that produces visually convincing synthetic images. However, their method does not produce corresponding ground-truth labels, and hence the data cannot be used as a benchmark, and evaluation remains limited to qualitative inspection. Their focus lies on data augmentation for training Content-Aware Image Restoration (CARE) [23] models, rather than on providing a validated dataset for systematic comparison.

Several generative models synthesize medical or biological images by conditioning on segmentation masks or other geometric annotations. For example, SPADE [24] introduces spatially adaptive normalization to generate high-fidelity images from semantic masks, and BrainSPADE [25] applies this idea to biomedical MRI synthesis, showing that mask-to-image generation can support downstream segmentation. Similarly, ControlNet [26] guides diffusion models using additional input images, such as edges or segmentation maps, to specify where objects appear and how they are arranged, through an auxiliary control branch. SegGuidedDiff [27] extends this to anatomically informed medical image generation by concatenating segmentation masks at every denoising step. Despite architectural differences, all of these methods depend on manually annotated segmentation masks to guide the image generation process. In contrast, our approach does not need ground-truth annotations: it extracts statistical structure directly from MT images and produces synthetic images with segmentation masks through a simulation-based pipeline, removing the dependency on manual segmentation entirely.

DRIFT [28] recently introduced a recurrent approach for instance segmentation of filamentous objects, motivated by the way humans “pick and trace” filaments. To enable training and evaluation, they generate synthetic datasets of curved lines with varying widths and lengths. Although this constitutes a creative strategy to alleviate data scarcity, the resulting images are highly abstract — essentially binary line drawings — and deviate strongly from the visual statistics of real microscopy data (see Fig. A.5b for examples). Their evaluation is centered on a custom recurrent model and comparisons to prior filament-tracing methods (SOAX [29], SIFNE [30], [31], and [5]), but the data and benchmark are not designed to capture the variability in appearance of actual MT microscopy images, nor to evaluate the performance of foundation models for segmentation. In contrast, our image pipeline explicitly aligns with real images in terms of model features (by design) and the biological plausibility of the resulting SynthMT dataset is judged by domain experts. Additionally, we make this data openly accessible via Hugging Face for easy benchmarking or usage for training.

### Generalist segmentation models

Recent work on foundation models for microscopy segmentation includes CellSAM [10], *µ*SAM [11], and Cellpose-SAM [12], all of which adapt SAM [32] to microscopy tasks. SAM has since been updated to SAM2 [33] and SAM3 [9], adding video segmentation and concept-specific text prompting. CellSAM employs a vision transformer to localize bounding boxes that are then passed to an adapted SAM module, removing the need for manual prompts. *µ*SAM adds a new decoder trained on microscopy data. Cellpose-SAM combines a modified SAM encoder with the flow field prediction and gradient tracking of Cellpose [34]. Furthermore, StarDist [18, 35] models are explicitly designed for segmenting star-shaped objects; although MTs are not star-convex, we include StarDist for completeness.

All of these models were benchmarked on a broad range of datasets [34, 36–59], most of which target nuclei or whole-cell segmentation. A few include elongated or tubular structures (e.g., filamentous bacteria in Omnipose [38], DeepBacs [39], or mitochondria in EM datasets), but none involve cytoskeletal MTs, where geometric continuity and sub-pixel-width features are critical.

Foundation models for general cellular and nuclear segmentation have reached a level of performance where further improvements are increasingly limited by annotation variability rather than model capacity. For example, Pachitariu et al. [12] report that Cellpose-SAM exceeds inter-annotator agreement on standard cell and nucleus benchmarks. In contrast, important challenges remain in settings like ours, where the objects of interest are elongated, filamentous structures, rather than compact objects like individual cells or nuclei.

### Microtubule-specific methods

Several earlier works have proposed custom pipelines for MT segmentation, and in some cases also tracking, often involving their own (synthetic) datasets. These include FIESTA [8], SOAX [29], SIFNE [30], MTrack [60], DRIFT [28], the methods by Liu et al. [5, 61, 62] and Masoudi et al. [63], KnotResolver [64], the work by Laydi et al. [19], and TARDIS [7]. Classical approaches typically depend on high signal-to-noise ratio (SNR) data and require substantial manual input, keeping humans firmly in the loop. Machine-learning-based methods [5, 7, 19, 28, 61–63] attempt to address this limitation; however, their accessibility is often hindered by a reliance on proprietary MATLAB software rather than open-source Python environments. Furthermore, the specialized datasets used to develop these methods are not released to the public.

As a result, reproducibility and fair cross-method benchmarking remain limited. Crucially, none of these approaches have been evaluated against modern foundation models such as those built on SAM [32]. Within the MT community, a notable exception in terms of availability and generalization capabilities is TARDIS [7]. It combines CNN-based segmentation with graph-based instance segmentation and is released as open-source Python code that provides pretrained models, including one specialized for MTs in 2D total internal reflection fluorescence (TIRF) images. The authors of TARDIS report that compared to Amira [65], a commercial software suite for 3D visualization and analysis, annotation accuracy for MTs improves by 42%.

In this work, we choose FIESTA as a traditional baseline and add TARDIS as a domain-specific foundation model. Our pipeline and evaluation framework are implemented in Python with open-source dependencies, enabling reproducible, cross-lab benchmarking while avoiding proprietary tools such as MATLAB (except for FIESTA).

We document all evaluated methods in Appendix A.1, detailing implementation choices and practical requirements. Beyond benchmarking, SynthMT is designed as a practical resource for experimental biologists, providing accessible, validated tools for MT analysis.

## 3 Mathematical framework for synthetic image generation

We formalize our image generation pipeline as a two-step stochastic process that generates synthetic microscopy images *I* ∼ *P_θ_*(*I*) conditioned on a parameter set *θ* as illustrated in Fig. 2. These two steps are: (1) a purely geometric process imitating microtubule (MT) morphology, and (2) an imaging process that adapts these structures such that they resemble real images. This second step is largely inspired by the augmentation pipeline used in AnyStar [17].

**Figure 2.**
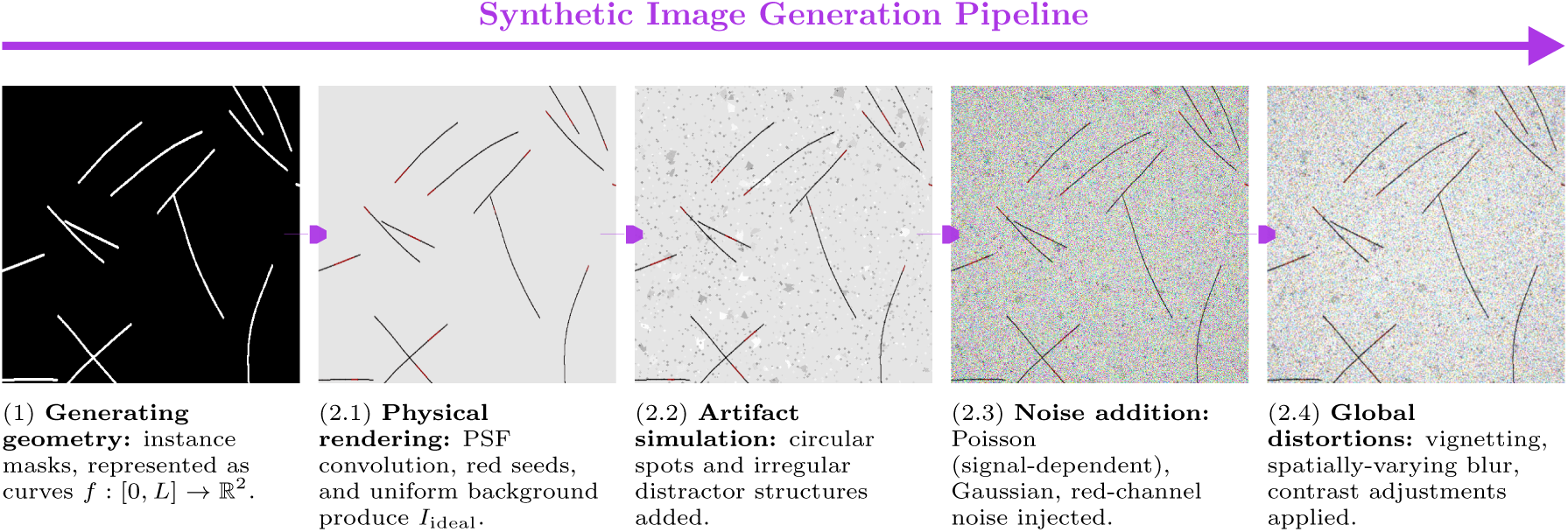
Our synthetic data generation pipeline produces realistic microtubule (MT) images with corresponding instance segmentation masks conditioned on a parameter set *θ*. (1) Generating geometry creates instance masks from geometric parameters (count, length, curvature) using polylines. (2.1) Physical rendering applies point-spread function (PSF) convolution to replicate optical properties, and adds red seeds and uniform background. (2.2) Artifact simulation introduces realistic distractor features (circular spots, irregular structures). (2.3) Noise addition models signal-dependent (Poisson) and signal-independent (Gaussian) noise sources. (2.4) Global distortions apply spatially-varying effects (vignetting, blur, contrast variations) to match real microscopy conditions. This approach enables the generation of labeled data that closely approximates experimental interference reflection microscopy (IRM) images, when its set of generation parameters *θ* is tuned accordingly (as explained in section 4).

**(1) Microtubule geometry.** Each MT is modeled as a polyline consisting of *n* segments 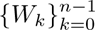. Together, these segments define a curve *f* : [0*, L*] → R^2^. Each *W_k_* is specified by a segment length *ℓ_k_* and a relative bend angle *ϕ_k_*, where *ℓ_k_* is sampled from a Gaussian distribution N (*µ, σ*^2^) with *µ, σ* ∈ *θ*. Filament curvature is introduced through a stochastic evolution of the bend angles *ϕ_k_*, which evolve as

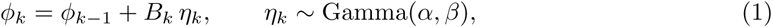

where *α, β* ∈ *θ* are shape and scale parameters, and *B_k_* is a random variable with values in {±1} that determines whether the filament bends to the left or right at segment *k*. The sampling of *B_k_* is determined through a flip probability *p*_flip_ ∈ *θ* and the maximum number of flips *n*_flips_ ∈ *θ*,

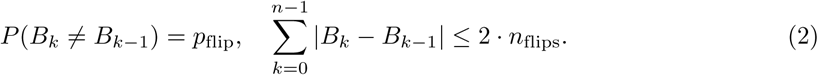

This yields a correlated random walk with persistence length governed by (*α, β*). The overall filament length *L* is drawn from a Gaussian distribution parametrized by *θ*, and the initial seed orientation *ϕ*_0_ is sampled uniformly from [0, 2*π*). Because segments are allowed to span only a few pixels, after step (2) they form smoothly curved filaments.

### Synthetic Image Generation Pipeline

**(2) Image rendering.** Given a curve *f* as constructed before (i.e., a piecewise-linear MT skeleton), we compute a binary mask *M_S_* of shape (*h, w*) ∈ N^2^ that marks all pixels intersected by *f* . This mask is then convolved with the point-spread function (PSF) *psf* , and scaled by a contrast factor *A* ∈ *θ* and background intensity *B* ∈ *θ*,

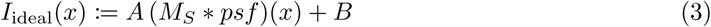

for every pixel *x* ∈ ([0*, h*) × [0*, w*)) ∩ N^2^. Through convolution with the PSF, intensity spreads to neighboring pixels, which determines the filament’s physical width in the rendered image.

Finally, noise and imaging artifacts are introduced via a stochastic operator *ν_θ_* that combines multiple perturbation processes. These include signal-independent noise (e.g., additive Gaussian noise), signal-dependent effects (e.g., Poisson shot noise and multiplicative speckle), channel-specific distortions, spatially correlated background variations, and structured artifacts such as vignetting, blur, and random distractor spots. All of these random variables instantiated by the noise process are collected in *θ* as well. This yields the final image *I* through

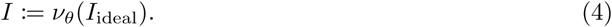

### Two-step stochastic process

Together, the complete pipeline defines a probability distribution *P_θ_*(*I*) over synthetic images, where *θ* encodes the distribution of the number of MTs, their length distribution and curvature statistics, and further imaging parameters. In principle, this framework also extends to a temporal process by defining a Markov chain {*I_t_*} (e.g., including MT dynamics such as growing, shrinking, pause with Gaussian growth/shrinkage increments at each time step). In the present work, however, we restrict to the *single-frame setting*, that is, single-sampling *I* ∼ *P_θ_* without simulating multi-frame temporal dynamics.

## 4 Methods

We describe the creation of SynthMT, the first dataset of annotated, synthetic *in vitro* microtubule (MT) microscopy images that closely mimic the visual characteristics of real experimental data.

Furthermore, we provide the protocol for assessing their perceptual realism using domain expert judgments and for evaluating fully automated segmentation methods on both SynthMT and a small labeled set of real images.

### Creating SynthMT and annotating real data

With the mathematical formulation of the image generation process in place (see section 3), we summarize the full pipeline to construct SynthMT, illustrated in Fig. 3: For a given reference distribution *Q* consisting of real, unlabeled images, the generator *P_θ_* iteratively creates images *I* ∼ *P_θ_*(*I*) that are compared to samples from *Q*. For this comparison, we embed the images using DINOv2 [66], a pre-trained state-of-the-art vision transformer, designed to generate rich visual features. The pipeline parameters *θ* are then tuned to minimize the resulting embedding distance, without requiring ground-truth annotations.

**Figure 3.**
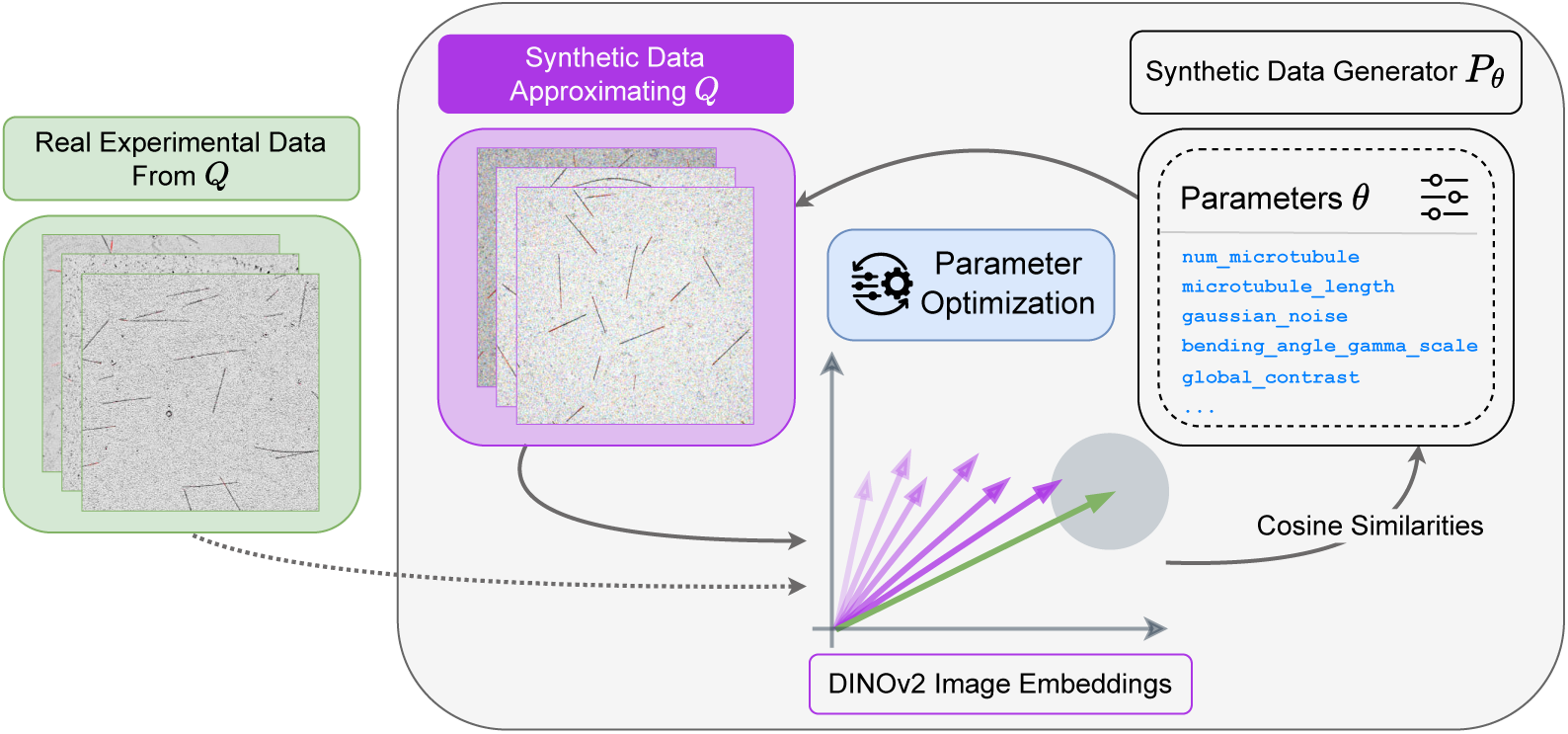
Optimizing *θ* aligns synthetic image distributions with real, annotation-free microscopy data. Real interference reflection microscopy (IRM) images (left) and synthetic images (center) are embedded using DINOv2. The parametric generator *P_θ_* (right) creates images by sampling from distributions governing geometric properties (filament count, length, curvature) and imaging characteristics (PSF, noise, artifacts, contrast, distortions), all controlled by *θ*. An optimization loop iteratively refines *θ* by maximizing cosine similarity between real and synthetic embeddings, ensuring that synthetic images match the statistical properties and visual characteristics of experimental data.

### Real data

The real images for our empirical target distributions *Q_i_* were provided by two wet labs that routinely investigate reconstituted MTs. In total, we obtained 44 real IRM videos, each containing between 30 and 360 frames, spanning multiple experimental conditions and microscope setups, and capturing substantial diversity in MT geometry and visual properties such as contrast, noise, pixel intensity distributions, and background textures. Four representative examples of this variability are shown in Figure 4. More information about this real data can be found in appendix A.3. These raw videos were histogram-normalized and background-subtracted^1^, with no additional processing applied.

**Figure 4.**
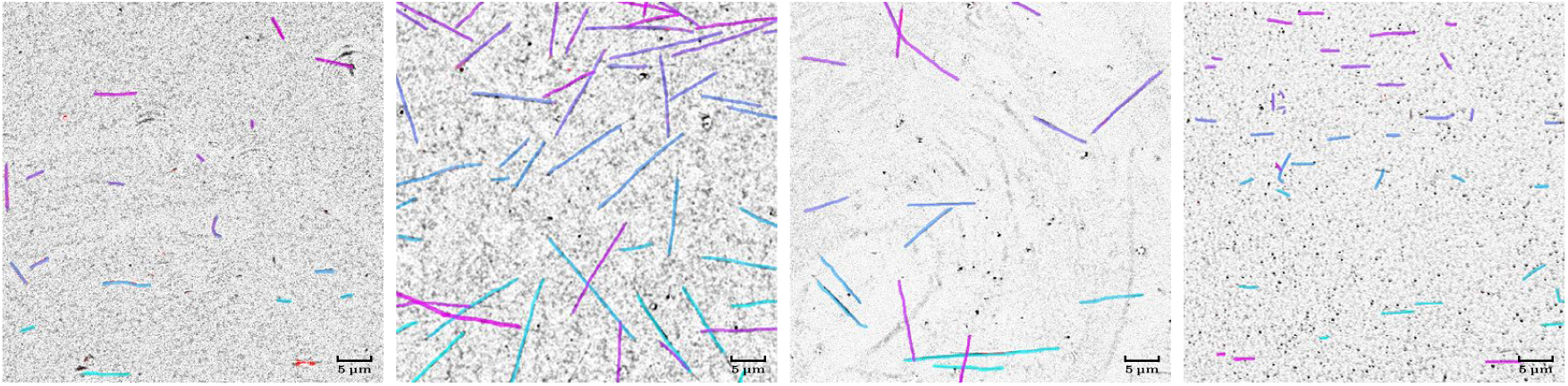
Real IRM images with human ground-truth labels (for evaluation only). Four exemplary frames from the 66 video crops of size 512 × 512 used to establish target distributions {*Q_i_*}^66^ (each containing 10 randomly sampled frames). Each image shows individual *in vitro* MTs growing from stabilized seeds under different experimental conditions, exhibiting natural variation in quantity, length, curvature, overlapping MTs, contrast, noise characteristics, filament density, and background properties. These target distributions act as references in the optimization process (Fig. 3), where DINOv2 embeddings of *Q_i_* guide the synthetic data generation. Ground-truth labels from one annotator are overlaid to illustrate the filament structures present in the real data; they are used solely for later method evaluation and are not required for generating the synthetic dataset SynthMT.

From these videos, we take random 512 × 512 pixel crops. We manually discard crops containing significant imaging artifacts or distortions that would typically be excluded from a standard analysis, yielding a final set of 66 smaller videos. Each cropped video is treated as a distinct target domain *Q_i_*, *i* = 1*, . . . ,* 66, from which we sample 10 random frames to serve as reference images for optimization. Importantly, for later method evaluations on unseen, real data, one of these frames from each crop is manually annotated by two independent human annotators using Label Studio [67]. In the analyses, we designate one annotator’s labels as ground truth and treat the other annotator’s labels as predictions, yielding an inter-annotator agreement score that quantifies the inherent variability of manual MT segmentation. This human baseline provides a reference point for assessing whether automated methods achieve human-level accuracy. Fig. 4 shows four exemplary frames together with the annotations used as ground truth.

### Optimizing *θ*

To align statistical and visual properties of the generated synthetic images with those of the selected real microscopy data {*Q_i_*}^66^ , we utilize the tree-structured Parzen estimator (TPE) [68] with 1000 iterations for each *i* = 1*, . . . ,* 66. TPE has repeatedly been shown to outperform alternative optimization algorithms [69, 70] and has become the de-facto standard in recent Hyperparameter Optimization (HPO) frameworks, where it is commonly used as the default sampling strategy [71, 72].

As a shared embedding space for comparing generated and real images, we choose the self-distillation method DINOv2 [66], which correlates well with human similarity judgments [73]. Moreover, as shown by Bolya et al. [74], the most informative embeddings for perceptual similarity are often obtained from intermediate rather than the final layers. This observation aligns with our findings: features from the fifth DINOv2 layer, denoted DINOv2_5_, balance abstraction and sensitivity to the subtle, fine-grained image structures that define IRM MT data. Altogether, the optimization maximizes the cosine similarity

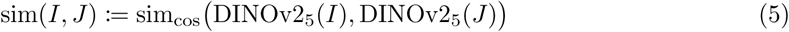

between generated images *I* ∼ *P_θ_*(*I*) and real samples *J* from *Q_i_* in the feature space. We aggregate the similarity between synthetic and real frames using the *maximum similarity* across the 10 references in *Q_i_* as the objective. This guarantees that the chosen parameters yield synthetic images that are plausible for at least one representative frame in *Q_i_*. Finally, we retain the top *k* = 10 candidate solutions 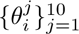 that yield the highest aggregated similarity. SynthMT is the product of drawing 10 images for each 1 ≤ *j* ≤ 10 and 1 ≤ *i* ≤ 66, yielding 6600 images of size 512 × 512, examples of which are shown in Fig. A.4. The masks are stored as 3D TIFF images, where each slice is the mask assigned to a single instance. This preserves overlapping objects correctly and resembles the output of SAM (see appendix A.1).

SynthMT is publicly hosted on Hugging Face under a stable organization account at huggingface.co/HTW-KI-Werkstatt/SynthMT. The dataset card provides versioning, update logs, and contact information, and we will maintain SynthMT by releasing documented updates if errors are discovered or extensions are added. Users can contact the corresponding authors to report issues, request corrections, or raise takedown concerns.

### Assessing the perceptual realism of SynthMT

We conduct a study with *n* = 6 domain experts to evaluate the quality and realism of our synthetic microscopy images. To enable a direct comparison, we include 10 real images, 10 synthetic images from SynthMT, and 10 images from the only openly available synthetic dataset for *in vitro* MTs, DRIFT [28]. All 30 images are presented in randomized order and each image is shown exactly once to each participant. The study is administered as a web-based interface (see appendix A.5 for details). Two of the six participants are co-authors of this paper; neither was involved in the data generation pipeline.

Experts rate each image on five aspects D using a 7-point Likert scale (1 = very poor, 7 = excellent): (a) Shape and general appearance of the MTs (structural fidelity), (b) Realism of the background, (c) Lighting, intensity, and blurring realism, (d) Noise pattern realism, and (e) Overall quality. The five aspects are motivated by quality evaluation criteria for microscopy imaging [75], while the use of aspect-specific Likert ratings follows recent generative evaluation studies emphasizing structured perceptual assessment of image realism [76, 77]. We use this setup rather than pairwise forced-choice comparisons [78–81] to obtain aspect-specific judgments that also inform future pipeline improvements.

### Distributional comparison of ratings

Experts differ systematically in their use of Likert scales (e.g., lenient vs. strict raters). To remove these fixed rater effects, we *z*-normalize each expert’s scores prior to aggregation. In particular, for expert *e* = 1*, . . . ,* 6 and aspect dimension *d* ∈ D, we compute the within-expert mean *µ_e,d_* and standard deviation *σ_e,d_* across all images, and transform raw ratings *r_e,d_* into normalized scores

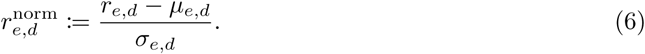

We visualize the normalized ratings for real and synthetic images using violin plots per dimension (Fig. 5), displaying the full rating density, median, and interquartile range. Because normalization is done per expert, these visualizations capture differences in perceived quality rather than rater-specific scale usage.

**Figure 5.**
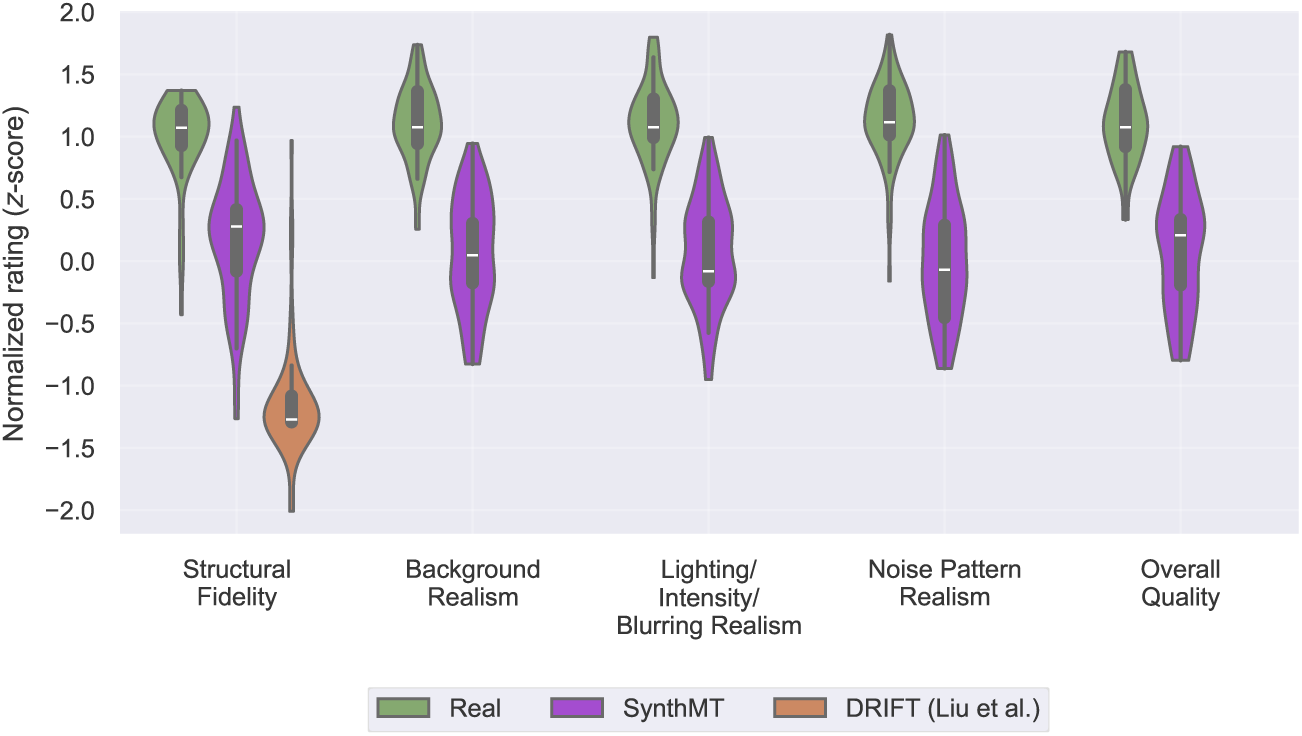
Domain experts confirm perceptual quality of SynthMT images. Violin plots show *z*-normalized ratings of *n* = 6 domain experts across five quality dimensions for real IRM images and synthetic images from SynthMT and DRIFT [28] (10 images each, 30 in total). Each violin displays the full distribution, median (white line) and interquartile range (thick bar). Ratings were collected on a 7-point Likert scale. DRIFT permits evaluation only of structural fidelity due to its black-and-white outputs (see exemplary images in Fig. A.5b). SynthMT images score higher than DRIFT on this dimension, indicating that parameter-optimized synthesis yields structures that more closely resemble real microscopy data. A measurable gap to real images persists across all dimensions. Nevertheless, experts rate the backgrounds, lighting, and noise patterns of SynthMT as internally coherent and plausibly aligned with real IRM, in contrast to DRIFT’s limited realism.

### Evaluations on SynthMT and real data

As discussed in the related work (section 2), we focus on the classical algorithm FIESTA [8] as a baseline, and compare to the foundation models SAM [32], SAM2 [33], *µ*SAM [11], CellSAM [10], Cellpose-SAM [12], and SAM3 [9] (listed chronologically by release date). StarDist [35] has an improper geometric bias for this task but is considered for completeness, and TARDIS [7] is added as the only MT-specific foundation model. The implementation and parameters of each method are detailed in appendix A.1. All methods are run in their “automated” mode without further input from our side. The only exceptions are SAM3Text, for which we use our manually specified default prompt “*thin line*”, and FIESTA, where we set the *full width at half maximum* (FWHM) parameter to 3. We ignore other methods that cannot work in a fully automated fashion or are not publicly available (e.g., Masoudi et al. [63] or MTrack [60]). Initial tests with a StarDist model trained on AnyStar [17] showed complete failure on our 2D data. The authors later verified^2^ that the method is inherently ineffective in this setting.

We evaluate two categories of metrics: segmentation quality and downstream biological performance, as detailed below.

### Preprocessing

We disable method-specific preprocessing steps and instead gather all of those in our own preprocessing function. For details, see appendices A.1 and A.2. This ensures that all methods share the same preprocessing capabilities during hyperparameter tuning (see HPO below).

### Method outputs

The methods differ in output format, which becomes crucial in postprocessing and metric computation. In particular, SAM, SAM2, and SAM3 return lists of 512 × 512 boolean masks, one per instance, which resembles the format of our labels for SynthMT and the real images we annotated. StarDist, *µ*SAM, CellSAM, and Cellpose-SAM produce 512 × 512 integer arrays, where 0 denotes background and each positive integer corresponds to a unique instance. We convert these into our format by splitting the integer-labeled arrays into separate boolean masks, one per nonzero label. FIESTA and TARDIS do not output segmentation masks as arrays. Instead, they return lists of anchor points (*p_k_*)*_k_*, one for each instance. Whenever we report instance lengths, for mask-based outputs *M* our proxy is the skeleton pixel count *L* = |*S*(*M* )|, where *S*(*M* ) is the thinned, one-pixel-wide binary skeleton. For anchor points (*p_k_*)*_k_*, we sum Euclidean distances, *L* = ∑_k_ ∥*p_k_*_+1_ − *p_k_*∥_2_.

### Postprocessing

To ensure comparability across methods, we also disable all method-specific postprocessing routines and apply a unified postprocessing step implemented by us as follows. Predicted instances are filtered by length and area (for anchor-point methods, only length is used), requiring them to fall within dataset-specific minimum and maximum values. In our benchmark, these thresholds are computed automatically from the ground-truth labels of SynthMT resp. the real data. In practical applications, however, domain experts typically know the plausible range of filament lengths and areas for their particular imaging setup, making such filtering a natural and interpretable step. By removing implausible tiny fragments and related artifacts, this unified postprocessing ensures more stable and meaningful comparisons across segmentation methods.

### Segmentation metrics

Accurately evaluating segmentation of thin, elongated MTs is non-trivial: classical pixel-overlap metrics such as Intersection over Union (IoU) over-penalize small transversal misalignments while under-representing errors in biologically relevant properties like filament length, continuity, or curvature. For MTs that are only a few pixels wide, a slight lateral shift or minor width disagreement can disproportionately reduce IoU, even if the traced centerline (and thus the inferred length) is essentially correct. To emphasize geometry rather than raw area agreement, we adopt the Skeleton Intersection over Union (SKIoU) [61] metric.

For anchor-point outputs (i.e., outputs from FIESTA and TARDIS), we compute a corresponding instance mask by fitting a spline with smoothing condition s=0^3^ through these points. We then represent this spline as a binary mask *M* to match the format of the other methods.

Given predicted and ground-truth instance masks *M*_pred_ and *M*_gt_, and again denoting *S*(*M* ) as the skeleton of a mask *M* , their SKIoU is defined as

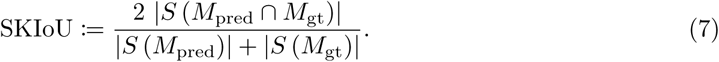

This provides a symmetric, length-normalized overlap over skeleton pixels: it attains 1 only when centerlines coincide, and it drops sharply when filaments are missing (under-segmentation), spuriously split or merged (topology errors), or substantially curved differently (shape distortion). Because skeleton extraction collapses width, SKIoU becomes robust to small diameter inconsistencies and lateral blurring artifacts that are common in IRM imaging.

For evaluation, we compute several common object detection and instance segmentation metrics based on the SKIoU values. We report the mean SKIoU of an image, calculated as follows. For each ground-truth instance, we find the best-matching prediction and record its SKIoU. If no matching prediction is found (a false negative), the score for that ground-truth instance is 0. The mean SKIoU value thus reflects segmentation quality while also penalizing for missed objects. While informative, this metric does not penalize false positives.

To provide a more complete picture, we also report Average Precision (AP) and F1 scores. AP is calculated as the mean of AP values at SKIoU thresholds from 0.5 to 0.95 in steps of 0.05, summarizing detection and segmentation accuracy. The F1 scores at specific SKIoU thresholds of 0.5 (F1@0.50) and 0.75 (F1@0.75) represent the harmonic mean of precision and recall. Comparing F1@0.50 and F1@0.75 allows us to gauge the consistency of a method’s predictions; a small drop-off between these values indicates that the method maintains high localization accuracy even at stricter evaluation criteria.

### Downstream biological metrics

Segmentation scores alone do not answer whether a method performs well at recovering biologically relevant filament statistics. To accommodate that, we report downstream biological metrics derived from instance geometries such as **counting** the number of predicted instances and gathering their computed **lengths** (as explained above).

Furthermore, we collect the average **curvature** of each predicted instance as follows: Anchor-point outputs are converted into masks by connecting the points via straight lines. Then, given any instance mask (directly from the method or converted from anchor points), we order skeletonized pixels by greedy nearest-neighbor and fit a parametric spline (*x*(*u*)*, y*(*u*)) with smoothing condition s=0 to preserve geometry. Finally, we compute the average curvature of this instance as the mean of the curvatures *κ*(*u*), where

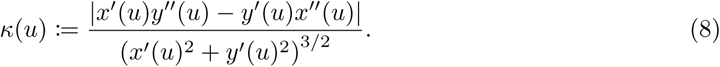

With these statistics at hand, we report mean and standard deviation across all images for counts, lengths, and curvatures. While this offers an intuitive understanding of systematic biases in the predictions relative to the ground-truth values, comparing distributions solely by their means and standard deviations is not sufficient. We therefore also compare the predicted and ground-truth distributions of length and curvature using normalized histograms with shared linear bins. Their dissimilarity is quantified through the Kullback–Leibler divergence (KL divergence) [82]

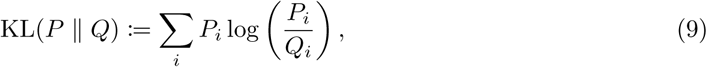

where *P* and *Q* denote the predicted and ground-truth discrete distributions, respectively (lower is better).

### Computational efficiency

We report computational throughput measured as processed images per second (“Img/s.” in Table 1). All results are obtained under sequential, unbatched inference to reflect per-sample efficiency. Experiments are conducted on a single NVIDIA A100 GPU; FIESTA is evaluated separately on a MacBook M3 Pro.

**Table 1.** Results on. SynthMT **signal unprecedented segmentation performance of the new SAM3 model.** For each method, we report segmentation metrics including Skeleton Intersection over Union (SKIoU) [61], Average Precision (AP) (mean over SKIoU thresholds), and F1 of SKIoU at 0.50 and 0.75. In addition, we report biological downstream metrics derived from counts per image, and length and curvature measurements per MT. These values are reported as mean ± standard deviation across all images and can be directly compared to the ground-truth statistics (last row). To capture distributional differences in greater detail, we further report the Kullback–Leibler divergence (KL divergence) between the predicted and ground-truth distributions of length and curvature. All methods are evaluated zero-shot using their default parameters. Rows marked with “+ HPO” show performance after Hyperparameter Optimization (HPO) maximizing SKIoU on 10 randomly sampled synthetic SynthMT images, illustrating few-shot adaptation potential. SAM-family models are run in Automatic Instance Segmentation (AIS) mode; SAM3Text uses SAM3’s text prompt mode (default prompt: “*thin line*”). For each column, the best-performing default and tuned methods are highlighted in **bold**, and ↑ (↓) signals higher (lower) is better.

| Model | Img/s ↑ | Segmentation Metrics | | | | Biological Downstream Metrics | | | | |
| --- | --- | --- | --- | --- | --- | --- | --- | --- | --- | --- |
| | | SKIoU ↑ | AP ↑ | F1 ↑ | | Count<br>*n*/img | Length | | Curvature | |
| | | | | @.50 | @.75 | | μm/MT | KL ↓ | μm<sup>-1</sup>/MT | KL ↓ |
| *Baseline* |  |  |  |  |  |  |  |  |  |  |
| FIESTA [8] | 0.21 | 0.12 | 0.12 | 0.22 | 0.18 | 99.63 ± 56.0 | 1.92 ± 4.3 | 5.03 | 6.74 ± 12.6 | 0.997 |
| + HPO | 0.16 | 0.24 | 0.25 | 0.39 | 0.30 | 38.80 ± 17.1 | 3.64 ± 5.9 | 3.74 | 6.90 ± 10.2 | 0.706 |
| *Foundation Models for Microscopy* |  |  |  |  |  |  |  |  |  |  |
| StarDist [35] | 4.93 | 0.01 | 0.04 | 0.00 | 0.00 | 1.85 ± 1.5 | 10.14 ± 2.9 | 0.69 | 7.67 ± 2.1 | 0.196 |
| + HPO | **8.64** | 0.31 | 0.34 | 0.36 | 0.15 | 20.36 ± 11.0 | 2.65 ± 1.4 | 1.95 | 7.79 ± 7.8 | 0.183 |
| TARDIS [7] | 0.32 | 0.45 | 0.48 | 0.59 | 0.45 | 17.15 ± 4.9 | 7.71 ± 6.5 | 0.56 | 8.28 ± 3.8 | **0.019** |
| + HPO | 0.34 | 0.48 | 0.55 | 0.57 | 0.41 | 12.33 ± 5.1 | 8.35 ± 6.3 | 0.41 | 8.50 ± 3.6 | 0.031 |
| μSAM [11] | **7.44** | 0.02 | 0.13 | 0.03 | 0.02 | 0.50 ± 1.0 | 6.03 ± 5.3 | 0.88 | 8.16 ± 8.1 | 0.130 |
| + HPO | 6.56 | 0.66 | 0.73 | 0.74 | 0.64 | 14.58 ± 22.9 | 7.22 ± 6.3 | 1.24 | 8.68 ± 10.5 | 0.132 |
| CellSAM [10] | 2.63 | 0.56 | 0.79 | 0.66 | 0.58 | 7.95 ± 4.6 | 9.57 ± 5.5 | 0.19 | 8.12 ± 4.1 | 0.021 |
| + HPO | 2.50 | 0.59 | 0.69 | 0.68 | 0.59 | **11.60 ± 6.7** | 9.33 ± 5.7 | 0.21 | 7.97 ± 4.5 | 0.031 |
| Cellpose-SAM [12] | 2.62 | 0.27 | 0.68 | 0.37 | 0.36 | 2.96 ± 3.2 | **12.54 ± 5.5** | 0.12 | **8.45 ± 4.2** | **0.019** |
| + HPO | 2.59 | 0.65 | 0.75 | 0.76 | 0.65 | 10.66 ± 6.0 | 10.39 ± 6.1 | 0.12 | **8.35 ± 4.9** | **0.012** |
| *General Purpose Foundation Models* |  |  |  |  |  |  |  |  |  |  |
| SAM [32] | 0.72 | 0.37 | 0.46 | 0.45 | 0.44 | 20.46 ± 17.3 | 3.91 ± 5.9 | 3.90 | 9.09 ± 22.5 | 0.700 |
| + HPO | 0.52 | 0.16 | 0.16 | 0.25 | 0.24 | 74.90 ± 37.8 | 2.01 ± 4.7 | 5.45 | 7.52 ± 18.4 | 0.912 |
| SAM2 [33] | 0.96 | 0.01 | 0.05 | 0.01 | 0.01 | 0.06 ± 0.3 | 3.65 ± 2.8 | 1.62 | 6.90 ± 4.1 | 0.269 |
| + HPO | 0.62 | 0.66 | 0.73 | 0.74 | 0.72 | 11.67 ± 5.3 | 11.89 ± 6.8 | 0.04 | 8.18 ± 4.1 | 0.014 |
| SAM3 [9] | 1.00 | 0.00 | 0.00 | 0.00 | 0.00 | 0.01 ± 0.1 | 3.02 ± 1.1 | 2.77 | 9.50 ± 3.9 | 0.489 |
| + HPO | 0.78 | 0.54 | 0.58 | 0.64 | 0.64 | 23.69 ± 26.8 | 6.64 ± 8.1 | 2.90 | 8.29 ± 13.6 | 0.364 |
| SAM3Text [9] | 4.26 | **0.85** | **0.86** | **0.91** | **0.90** | **13.03 ± 6.5** | 11.74 ± 7.1 | **0.07** | 7.54 ± 3.3 | 0.063 |
| + HPO | 3.97 | **0.93** | **0.95** | **0.97** | **0.96** | 11.73 ± 5.6 | **13.19 ± 6.7** | **0.02** | 7.56 ± 3.0 | 0.069 |
| Ground Truth | - | 1 | 1 | 1 | 1 | 11.28 ± 5.1 | 13.35 ± 6.7 | 0 | 8.42 ± 3.9 | 0 |

### Zero-shot setting

We evaluate all methods with their default parameters (as detailed in appendix A.1) on SynthMT and the unseen, real data.

### HPO-based few-shot setting

Furthermore, we automatically adapt both method and preprocessing hyperparameters using only 10 random samples from SynthMT. We conduct this HPO via the tree-structured Parzen estimator [68] with 1000 trials to maximize mean SKIoU. We report the resulting optimal configurations in appendices A.1 and A.2, and discuss optimization trajectories and parameter importance in appendix A.7.

## 5 Results

The resulting SynthMT dataset, as described in section 4, contains 6600 synthetic images with a varying number and shape of microtubules (MTs), and diverse backgrounds and noise. They are accompanied with segmentation masks for every MT present in each image. Moreover, two independent human annotators created instance segmentations for 66 real interference reflection microscopy (IRM) images.

### Perceptual realism of SynthMT

The images in SynthMT were constructed to match real IRM-like data in the fifth layer representation of the DINOv2 model (see Eq. (5)). For adoption of this dataset within the community, perceptual adequacy to humans matters as well. Here, we present the results of the expert study outlined in section 4.

While a gap relative to real images remains, the distributions in Fig. 5 nevertheless demonstrate that SynthMT occupies an intermediate but substantive region between DRIFT [28] and real data: SynthMT is geometrically closer to the real domain, perceptually coherent across background, lighting, and noise, and stable across experts. This establishes SynthMT as the most realistic synthetic alternative currently available.

In the following sections, we show that this perceptual adequacy has functional consequences too: methods’ strengths on SynthMT transfer to unseen, real IRM data, and tuning via Hyperparameter Optimization (HPO) on only 10 images from SynthMT further improves performance on both synthetic and real data for many methods.

### Benchmarking SynthMT and real data

Beyond our generation pipeline used to produce SynthMT, Table 1 depicts another main contribution of this work: benchmark results for all nine introduced methods across the 6600 synthetic images contained in SynthMT. Additionally, we present results on unseen, real data in Table 2, and a qualitative analysis in Fig. 8. The benchmark is fully reproducible with the code available at github.com/ml-lab-htw/SynthMT.

**Table 2.**
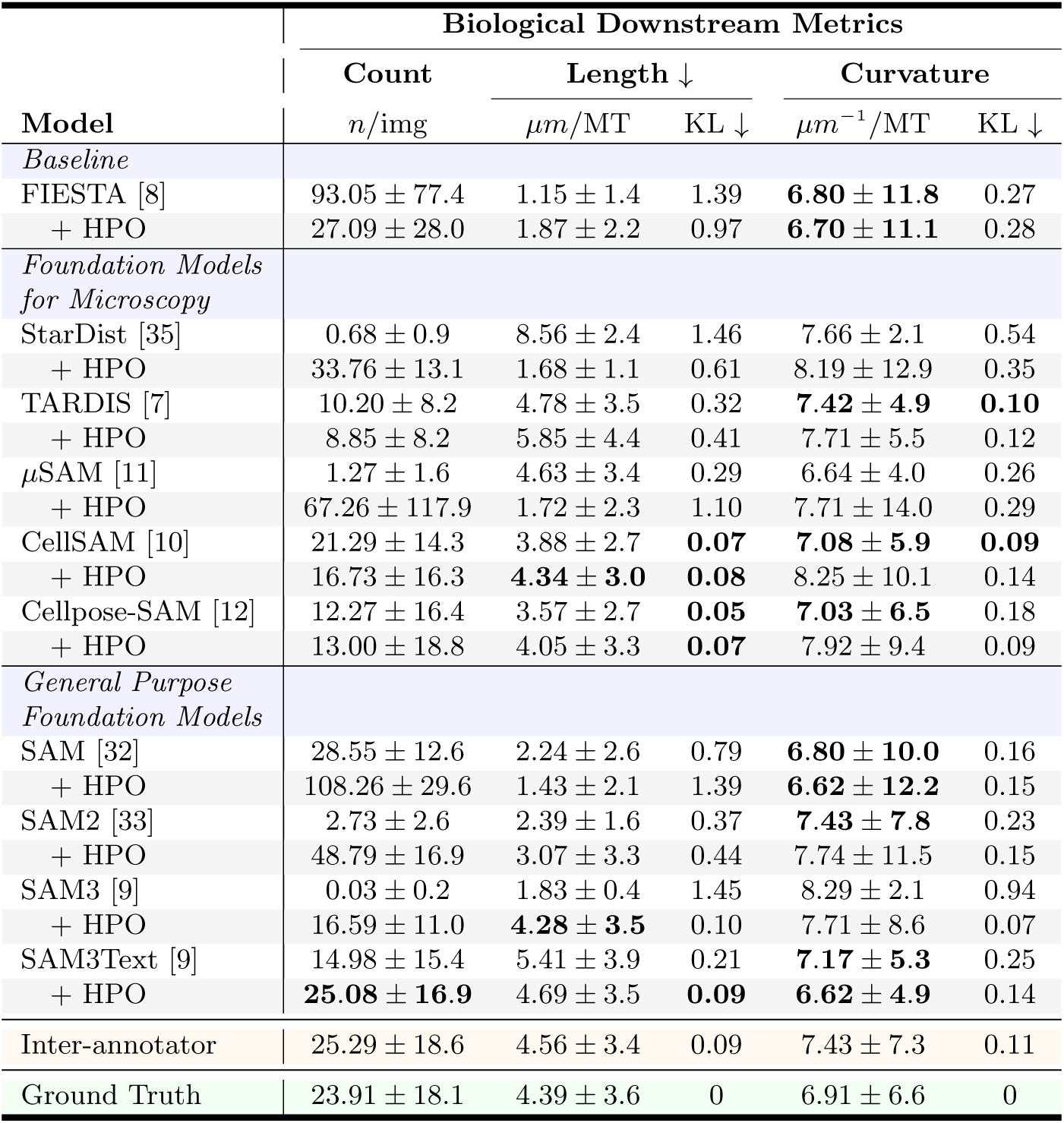
SAM3 signals capabilities of fully automated MT analysis on unseen, real IRM data. Analogous to Table 1, we evaluate methods on biological downstream metrics. Additionally, we report an inter-annotator baseline by treating the annotations of a second, independent human annotator as predictions and comparing them against the ground-truth annotations of the first annotator, thereby quantifying inherent variability in manual segmentation. The strategy of adapting hyperparameters of SAM3Text through an HPO on 10 randomly sampled SynthMT images leads to human-grade accuracy. Models that are on par with or surpass the human inter-annotator baseline are listed in **bold**.

### From FIESTA to SAM3Text

From Table 1 we find that all foundation models perform substantially better than the traditional, deep learning-free baseline FIESTA [8] which is still state-of-the-art in many labs. As one of the fastest methods, SAM3Text with tuned hyperparameters emerges as the candidate of choice for almost all tasks. Notably, on SynthMT it is the only evaluated method that achieves sub-micrometer accuracy in MT length estimation. Fig. 6 visualizes the close alignment between its predicted and ground-truth distributions of MT lengths and curvatures across scales. The only task where SAM3Text slightly lags behind is curvature estimation.

**Figure 6.**
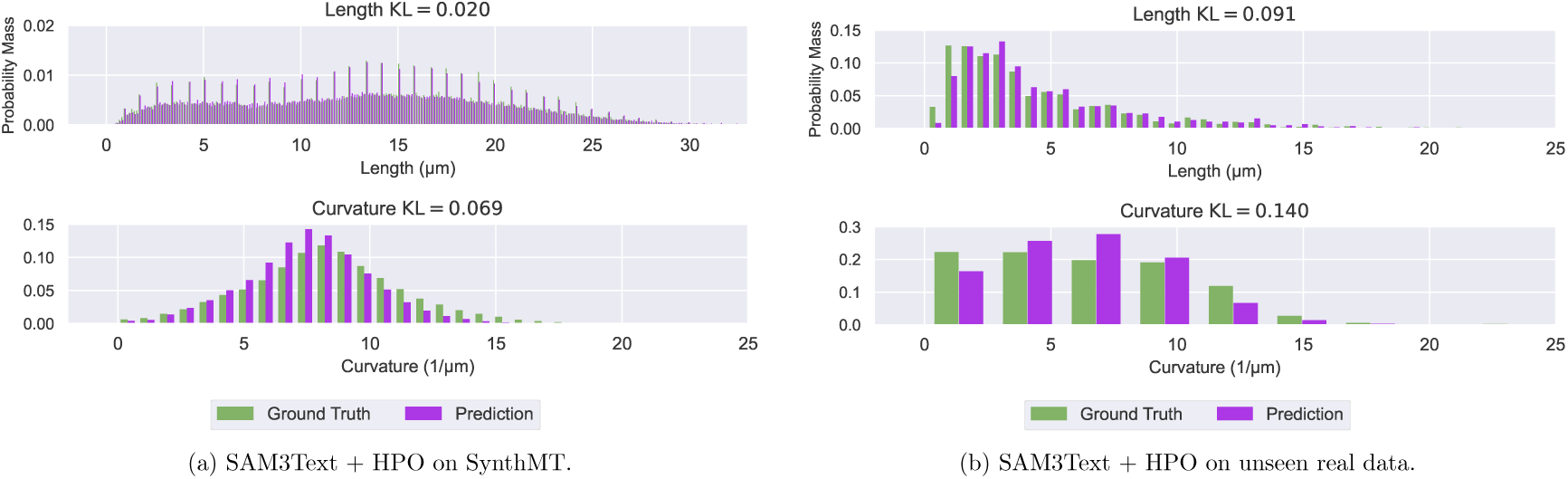
SAM3Text + HPO closely matches ground-truth MT length and curvature distributions across scales and datasets. Normalized histograms compare predicted and ground-truth distributions for MT length (top row) and curvature (bottom row) on (a) SynthMT and (b) unseen, real data. SAM3Text + HPO preserves both low and high values across the full range of lengths and curvatures, as reflected by low KL divergence values computed from these histograms. Distributions for the other methods on SynthMT are shown in Figs. A.8 and A.9.

### Improvements through HPO

In the few-shot (HPO) setting, we optimized method hyperparameters by maximizing SKIoU using only 10 random images from SynthMT. We find that this often greatly improves segmentation, except for TARDIS and CellSAM, which seem rather agnostic to parameters, and SAM, which shows a substantial drop in performance, indicating overfitting.

### Localization robustness

SAM3Text not only achieves the strongest segmentation scores overall, but also stands out for its internal consistency across complementary metrics. Its simultaneously high AP, SKIoU, F1@0.50, and F1@0.75 scores indicate that detected MTs are both reliably found and accurately localized, with little degradation when stricter overlap requirements are imposed. In particular, the stable performance from F1@0.50 to F1@0.75 suggests that SAM3Text predictions are geometrically precise. Other models from the SAM family exhibit a similar robustness pattern, albeit at a lower absolute performance level. In contrast, StarDist and TARDIS show a pronounced drop at F1@0.75, revealing inconsistencies between detection and precise alignment. A different failure mode is observed for CellSAM and Cellpose-SAM, which achieve comparatively high AP values while lagging behind in SKIoU. This discrepancy indicates that, although many MTs are detected, their spatial alignment is often imprecise, resulting in poorly localized segmentations. Such behavior is consistent with a higher incidence of missed or fragmented filaments in the later qualitative inspections.

### Segmentation metrics as proxies for biological downstream performance

Overall, improvements in segmentation metrics translate into better performance on biological downstream tasks. However, this relationship breaks down for methods whose segmentation quality is fundamentally insufficient. For methods with near-zero segmentation scores, such as StarDist, *µ*SAM, SAM2, and SAM3, even large gains in segmentation metrics achieved through HPO do not lead to meaningful improvements in biological performance and in some cases even degrade it.

Apart from that, *µ*SAM shows that poor performance on segmentation metrics is not immediately an indicator of bad downstream results. We hypothesize that this is because the default versions of SAM2, SAM3, Cellpose-SAM, and *µ*SAM did not generate any predictions on up to 60% of the 6600 images, while their HPO versions did for almost all. Hence, *µ*SAM identified only a small fraction of all MTs, yet statistically reproduced the correct distributions of MT geometries. We refer to Figs. A.8 and A.9 for these distributions for all methods, illustrating the curvature distribution match of *µ*SAM, even with poor predictions for count and length. Still, relying on downstream biological measurements when segmentation is weak comes at a risk, as they may be computed on noisy or incorrectly identified structures.

### Models made specifically for microscopy are strong on this task

Although SAM3Text consistently performs well, it is important to note that Cellpose-SAM, as a foundation model specifically released for microscopy tasks, also demonstrates strong performance on downstream metrics; even in its default setting (where it does not find any instances in 27% of the images). While TARDIS, *µ*SAM, and CellSAM lag behind (in line with the findings of Pachitariu et al. [12]), they easily beat generalist models such as SAM or SAM2 on these biologically relevant tasks. As expected, StarDist, although one of the fastest with ≈ 5 images per second, performs the worst with its default settings, since it is architecturally made for star-convex shapes. However, after tuning its hyperparameters it still outperforms FIESTA.

### Tests on unseen, real IRM data with human baseline further confirm SAM3Text

Analyzing performances on unseen, real data and comparing them to a human baseline allows us to put the previously presented numbers on the synthetic SynthMT dataset into context.

Fig. 7 demonstrates that SAM3, provided with a simple text prompt (“*thin line*”) and with optimized hyperparameters using 10 images from our synthetic dataset SynthMT, maintains strong performance on unseen, real images and achieves segmentation quality comparable to human performance. It reaches a mean SKIoU of 0.74 ± 0.18, compared to a human of 0.8 ± 0.14, placing the model’s performance within the variability observed between human annotators. Although other methods also benefit from HPO, none reaches this level of performance. The methods that do not improve through HPO are the same as those found above in the analysis on SynthMT (TARDIS,

**Figure 7.**
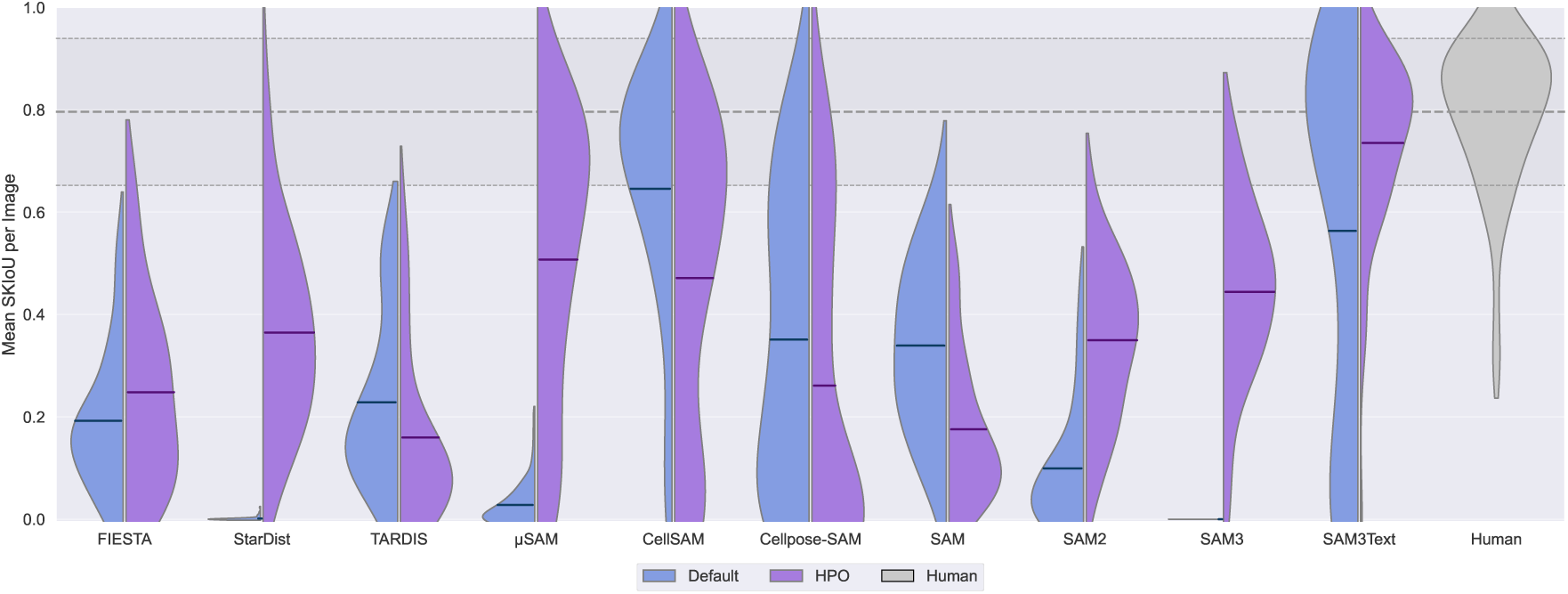
Only SAM3Text + HPO reaches human segmentation performance on unseen, real IRM data. Split violin plots show the distribution of per-image SKIoU scores (*n* = 66) for each method. The left side of each violin (blue) represents the default configurations, while the right side (purple) shows the performance after HPO on 10 random images from SynthMT. Each violin includes a horizontal line indicating the mean SKIoU across all images. The human performance is shown as a solid gray violin for reference. Its mean and standard deviation values are indicated by horizontal lines across the plot. The plot shows that the optimized SAM3Text matches the human inter-annotator baseline. While other methods also improve with HPO, none demonstrates the top-tier performance of SAM3Text. Notably, CellSAM already approaches human-level performance in its default configuration, but exhibits decreased performance after HPO.

**Figure 8.**
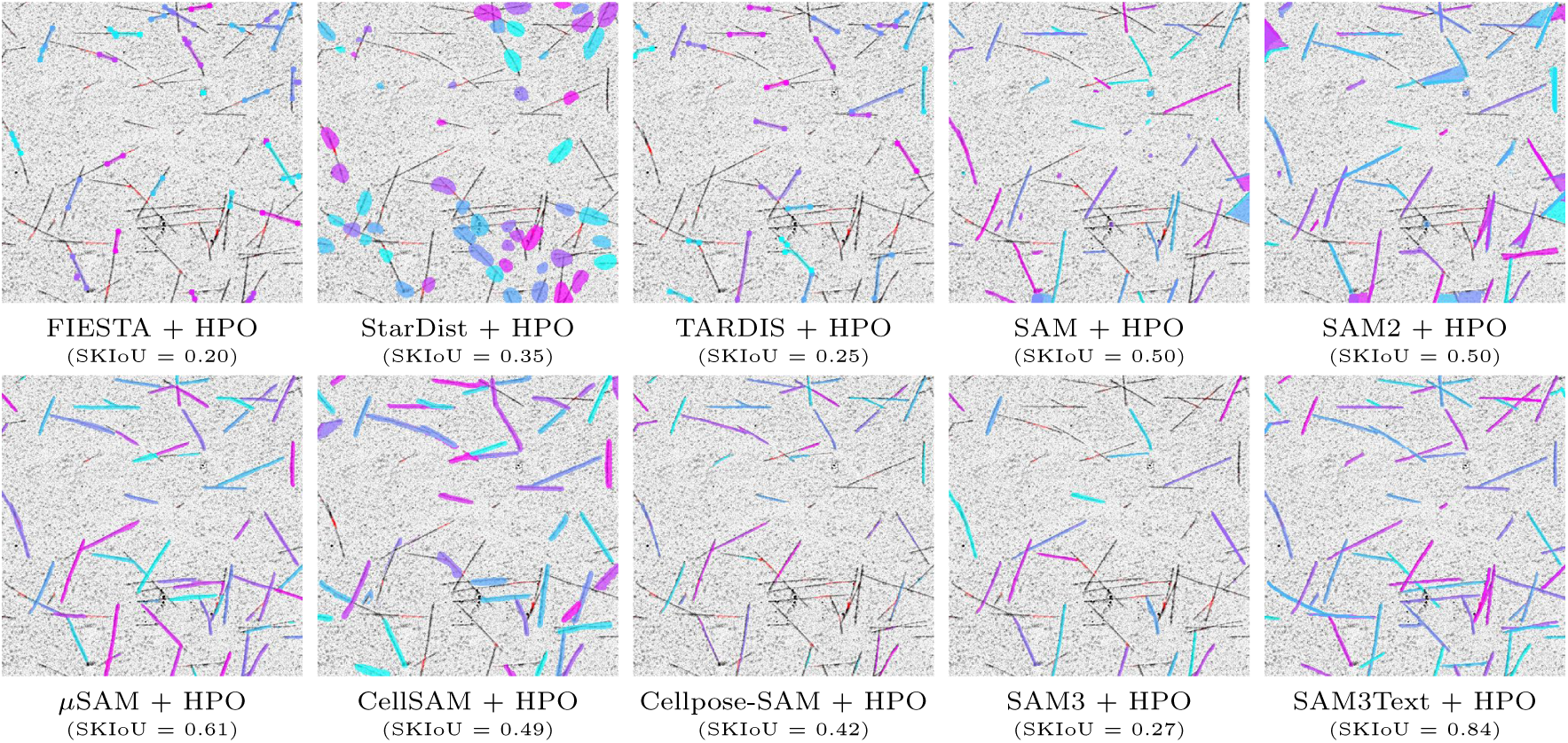
Qualitative comparison on an unseen, real-world *in vitro* reconstituted MT assay. For each method, we show predictions after HPO on 10 synthetic images from SynthMT (few-shot setting). The selected real image is particularly challenging, as it contains many intersecting MTs and exhibits a low signal-to-noise ratio (SNR), exposing a wide range of failure modes. For anchor-point methods such as FIESTA and TARDIS, only the first and last predicted points per instance are shown for visual clarity. Underneath each image we report the mean SKIoU value for this specific image, in order to correlate it with a visual impression. SAM3Text clearly performs best in this setting, while all other methods show limitations that may hinder their suitability for large-scale fully automated analysis. For more comparisons and dynamic exploration of this kind, we refer to our project page at DATEXIS.github.io/SynthMT-project-page.

CellSAM, SAM). Cellpose-SAM is the only method that improves on synthetic data, but worsens on unseen, real data.

Table 2 confirms these findings through the quantitative biological metrics, where SAM3Text consistently improves with HPO. It matches and slightly surpasses human inter-annotator agreement in counting accuracy, with a deviation of only 1.17 instances per image relative to the Ground Truth (GT), and achieves sub-micrometer accuracy in length estimation (deviation of 0.3 *µm*). For curvature, SAM3Text remains within the inter-annotator range, achieving a better average (0.29 *µm^−^*^1^ deviation) and only a marginally higher KL divergence.

However, all of the foundation models for microscopy — except StarDist — achieve reasonable downstream performance. In particular, CellSAM already performs strongly in its default configuration, ranking second in SKIoU. It also shows good agreement with the ground truth for length and curvature, but underperforms in counting accuracy. In contrast, only SAM3Text consistently attains top or near-top performance across all biological metrics without such compromises.

### Qualitative analysis reinforces the strength of SAM3Text

We examine a particularly challenging real example with many intersecting MTs and low SNR in Fig. 8 to qualitatively compare methods, and find that these observations are in line with the quantitative results reported before.

Among all methods tuned with HPO, SAM3Text clearly stands out. It consistently captures all visible MTs (see bottom right in Fig. 8), including most of the complex intersections, producing continuous and well-separated instances with only very few false positive or false negative detections.

Its predecessors exhibit diverse failure modes: SAM generates numerous small, spurious detections despite postprocessing, while SAM2 merges intersecting filaments and occasionally segments enclosed areas, producing incorrect, non-elongated shapes.

The anchor-point-based methods FIESTA and TARDIS, which were explicitly designed for MT analysis, show mixed performance. Both methods can resolve intersections, but they miss a substantial number of filaments. Moreover, when instances are detected, endpoints are often not identified correctly, leading to inaccurate length estimates.

Among the other foundation models specialized for microscopy, *µ*SAM, CellSAM, and Cellpose-SAM recover many filament structures but exhibit notable weaknesses. *µ*SAM and CellSAM frequently merge intersecting MTs, resulting in under-segmentation that hinders downstream measurements. CellSAM additionally produces overly thick segmentations. It is worth recalling that our SKIoU metric is robust to such width inaccuracies by design, as it operates on skeletonized versions of the predictions. Cellpose-SAM performs best within this group, correctly identifying a large fraction of instances, but still misses a large number of filaments, which limits its suitability for fully automated analysis pipelines that require high instance recall. Finally, StarDist fails to capture the elongated morphology of MTs as expected, since its star-convex object prior biases predictions toward circular shapes.

## 6 Discussion

### Suitability of automated microtubule (MT) analysis

A central question in MT image analysis is whether fully automated, expert-level segmentation is currently achievable. Our results provide a nuanced, previously unknown answer. Most segmentation approaches still fall short of the accuracy required for reliable downstream biological measurements on interference reflection microscopy (IRM) data of *in vitro* reconstituted MTs. They frequently fragment filaments, miss faint or short instances completely, or introduce spurious detections, even when Hyperparameter Optimization (HPO) is used (few-shot setting).

### SAM3 altered the picture

However, our study highlights a key positive outcome: guiding the recently introduced SAM3 model by a simple text prompt (“thin line”) and tuning it on only 10 synthetic SynthMT images reaches performance within the human inter-annotator range on unseen, real data. This is, to our knowledge, the first demonstration that a general-purpose vision foundation model can be adapted to deliver fully automated, biologically meaningful MT segmentation.

### Tuning with SynthMT enables human-grade performance

A key enabler of this achievement is the synthetic dataset SynthMT, which can be generated entirely from real MT images without the need for manual annotations. It automatically captures the visual statistics of real IRM frames and uses these to simulate filamentous structures. This enables the adaptation of segmentation methods to new microscopes, imaging conditions, or laboratories without any annotation effort. The success of SAM3Text after HPO on only 10 of those images demonstrates that method configuration can be effectively guided toward human-grade performance, without requiring any manual annotation of real data.

Within this context, SynthMT serves as both a benchmark and a diagnostic tool. It exposes the typical failure modes of existing methods, including fragmentation, missed instances, and sensitivity to domain shifts, and it provides a controlled setting for evaluating and tuning segmentation methods.

### Summary

SynthMT provides a reproducible and annotation-free framework for evaluating segmentation methods, identifying their limitations, and enabling powerful foundation models like SAM3 to be configured for filamentous microscopy data. This establishes a practical and fully automated route toward scalable MT segmentation suitable for high-throughput biological experiments, such as (non-dynamic) end-point experiments.

### Limitations and future work

One limitation is that our evaluation focuses exclusively on fully automated segmentation methods. Semi-automated approaches that require human input or corrections are not considered.

Moreover, our synthetic image generation relies on a parametric approach derived from real MT images. While it avoids the need for manual annotation, alternative approaches such as diffusion-based models have shown potential for generating realistic microscopy images [27], but they typically require large annotated datasets for training. Investigating these methods as a complementary or alternative generation strategy is an avenue for future work.

Furthermore, we did not use SynthMT as a large-scale training dataset for finetuning pretrained foundation models beyond the few-shot HPO experiments. SynthMT could potentially support fully supervised or semi-supervised training to further improve model performance.

Lastly, SynthMT currently focuses on 2D image generation from single static frames, which does not capture the temporal dynamics of MTs such as their alternating phases of growth and shrinkage. Extending the pipeline to videos would allow instance tracking over time, which is crucial for studying MT kinematics. Several approaches could be explored to achieve robust MT tracking. Time could be treated as a third dimension in volumetric models such as AnyStar [17] or Cellpose-SAM [12], or fully automated segmentation outputs could be combined with existing tracking algorithms [6, 60, 64]. Notably, SAM2 and SAM3 already support video data, which opens the possibility of extending their promptable, fully automated segmentation capabilities to temporal MT datasets.

## Acknowledgments

Our work is funded by the Deutsche Forschungsgemeinschaft (DFG, German Research Foundation) Project-ID 528483508 - FIP 12. We would like to thank Dominik Fachet and Gil Henkin from the Reber lab for providing data, and also thank the further study participants Moritz Becker, Nathaniel Boateng, and Miguel Aguilar. The Reber lab thanks staff at the Advanced Medical Bioimaging Core Facility (Charité, Berlin) for imaging support and the Max Planck Society for funding. Furthermore, we thank Kristian Hildebrand and Chaitanya A. Athale (IISER Pune, India) and his lab for helpful discussions.

## Author contributions

**MK** and **JW**: Conceptualization, Data Curation, Formal Analysis, Investigation, Methodology, Project Administration, Resources, Software, Validation, Visualization, Writing – Original Draft Preparation, Writing – Review & Editing.

**DF**: Data Curation, Resources, Writing – Original Draft Preparation.

**SR**, **FG** and **ER**: Conceptualization, Funding Acquisition, Supervision, Writing – Review & Editing.

## A Appendix

### A.1#Details about evaluated method

We summarize implementation and parameter details for all evaluated methods (ordered chronologically in terms of publication date): FIESTA [8], StarDist [35], TARDIS [7], SAM [32], SAM2 [33], *µ*SAM [11], CellSAM [10], Cellpose-SAM [12], SAM3 [9] (with Automatic Instance Segmentation (AIS)) and SAM3Text (SAM3 guided by a text prompt). StarDist, *µ*SAM, CellSAM, and Cellpose-SAM produce 512 × 512 integer arrays, where 0 denotes background and each positive integer corresponds to a unique instance. As each pixel can belong to only exactly one instance or background, overlaps cannot be represented. On the other hand, SAM, SAM2, SAM3 and SAM3Text return lists of 512 × 512 boolean masks, one per instance. This representation allows for overlapping instances, since multiple masks can contain the same pixel as foreground (instance). FIESTA and TARDIS do not output segmentation masks as arrays. Instead, they return lists of anchor points (*p_k_*)*_k_* = (*x_k_, y_k_*)*_k_*, one for each instance. For more information about the Hyperparameter Optimization (HPO) that yields the below stated “tuned” hyperparameter values, we refer to appendix A.7.

The full code, including fixed versions for full reproducibility, is available at github.com/ml-lab-htw/SynthMT.

#### FIESTA

Fluorescence Image Evaluation Software for Tracking and Analysis (FIESTA [8]) is a classical, non–machine-learning software tool for tracking fluorescently labeled filaments in 2D or 3D time-lapse microscopy data. We use FIESTA as a baseline, because it represents the traditional, widely adopted approach for microtubule (MT) analysis.

The tracking algorithm evaluates every image in a sequence independently before linking detected objects into trajectories. This involves several steps: thresholding, feature detection, image segmentation, a fitting process using Gaussian models to achieve sub-pixel localization, and interpolation. Finally, a graph-theoretic approach is used to link the detected objects into trajectories. The primary input is a time-series of images, and the output consists of data files with filament coordinates (anchor points).

In this work, we use FIESTA via a MATLAB implementation^4^ invoked from Python using the MATLAB Engine API for Python^5^. We utilize only the single-frame detection capabilities of FIESTA to identify filaments, without using its temporal tracking functionality. The fwhm_estimate parameter has no default value, so we fixed an estimate of 3.0 based on our data. Furthermore, we made several adaptations to the original FIESTA source code to ensure smooth execution, which we attribute to MATLAB version incompatibilities or our use of FIESTA in a scripted, non-GUI environment. The default and tuned hyperparameters are summarized in Table A.1.

**Table A.1.**
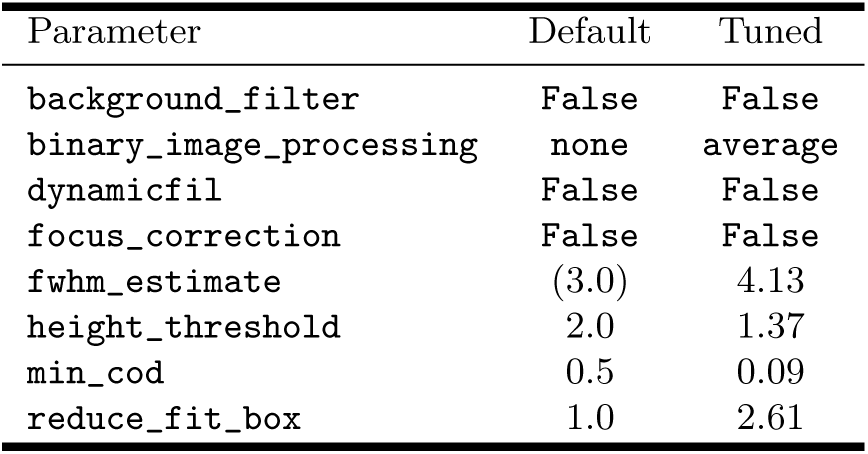
Optimized FIESTA hyperparameters obtained through HPO.

#### StarDist

StarDist is a deep learning-based object detection and segmentation method, particularly effective for identifying cell nuclei in microscopy images in densely packed scenarios. Instead of traditional bounding boxes, it represents objects as star-convex polygons, which are well-suited for the roundish shape of nuclei. The core of StarDist is a convolutional neural network (CNN) based on the U-Net architecture, that, for every pixel, predicts an object probability and the radial distances to the object’s boundary representing a star-convex polygon. The latter are finally refined using non-maximum suppression (NMS).

StarDist is used via model = stardist.models.StarDist2D.from_pretrained(pretrained)^6^, where pretrained can be one of 2D_versatile_fluo, 2D_versatile_he, or 2D_paper_dsb2018. Predictions on an image are called as

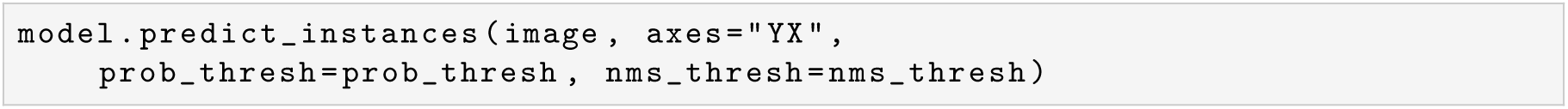

The default and tuned hyperparameters are summarized in Table A.2.

**Table A.2.** Default StarDist hyperparameters and those obtained through HPO.

| Parameter | Default | Tuned |
| --- | --- | --- |
| `pretrained` | `2D_versatile_fluo` | `2D_paper_dsb2018` |
| `prob_thresh` | 0.479 | 0.61 |
| `nms_thresh` | 0.3 | 0.03 |

#### TARDIS

Transformer and Rapid Dimensionless Instance Segmentation (TARDIS) is a fully automated segmentation workflow designed for cytoskeletal filaments and organelles. The TARDIS workflow consists of three main steps: semantic segmentation which produces a semantic mask, post-processing of the semantic mask into a point cloud representation of the objects, and instance segmentation of the point cloud. It is used via tardis_em^7^. Predictions on an image are generated by first saving it as a grayscale 2D TIFF in a temporary working directory work_dir and then calling

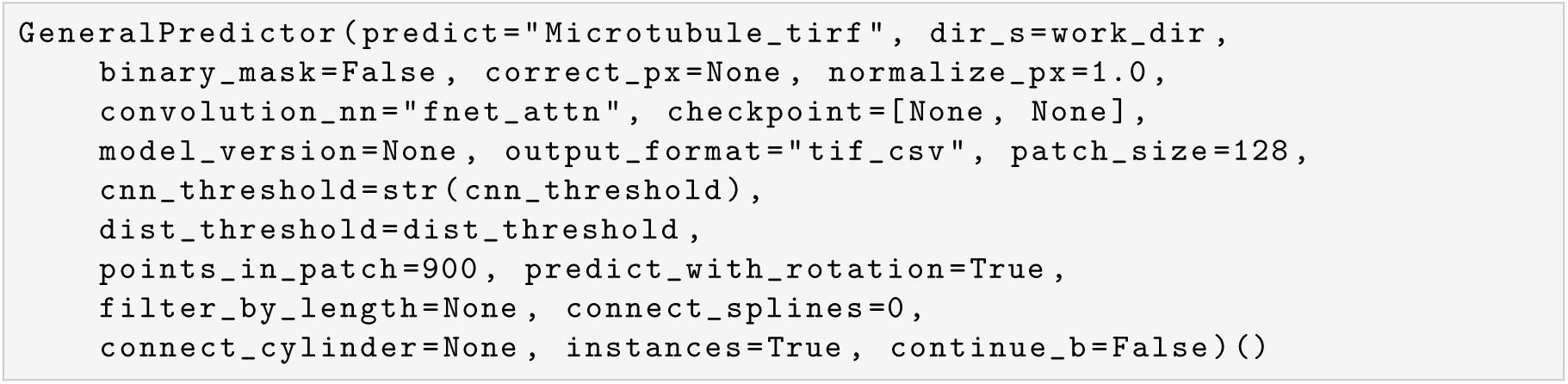

The default and tuned hyperparameters are summarized in Table A.3. Note that the predictions are saved as CSV files in work_dir from which the anchor points can be read.

**Table A.3.** Default TARDIS hyperparameters and those obtained through HPO.

| Parameter | Default | Tuned |
| --- | --- | --- |
| `cnn_threshold` | 0.1 | 0.18 |
| `dist_threshold` | 0.5 | 0.69 |

#### SAM

Segment Anything Model (SAM) is a foundation model for image segmentation. Trained on over a billion masks, SAM enables zero-shot transfer to new image distributions and segmentation tasks. We use SAM in Automatic Instance Segmentation (AIS) mode, which generates masks without manual prompts by sampling points across the image.

Its architecture features an image encoder, a prompt encoder for point or bounding box inputs, and a mask decoder. While the original paper also mentions text prompts, this feature was never made publicly available. We use SAM via the Hugging Face transformers.pipeline^8^ wrapper.

Instantiation happens via model=pipeline(“mask-generation”, model=“facebook/sam-vit-huge”) and prediction on an image are called via

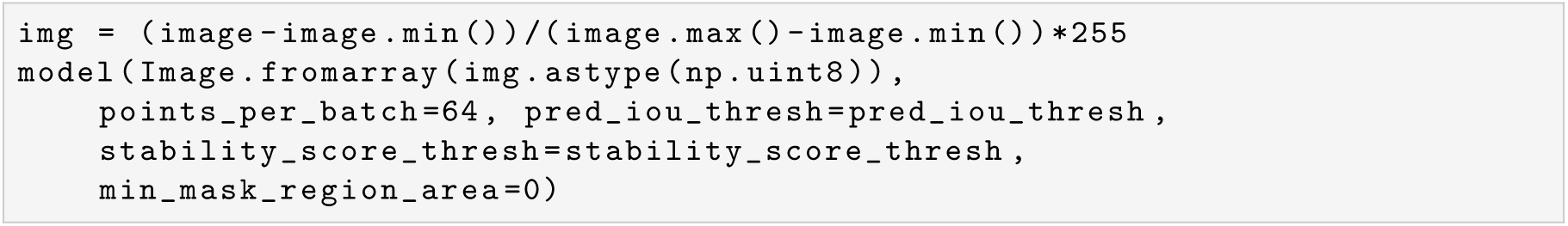

The default and tuned hyperparameters are summarized in Table A.4.

**Table A.4.** Default SAM hyperparameters and those obtained through HPO.

| Parameter | Default | Tuned |
| --- | --- | --- |
| `pred_iou_thresh` | 0.88 | 0.56 |
| `stability_score_thresh` | 0.95 | 0.93 |

#### SAM2

Segment Anything Model 2 (SAM2) is a unified model for both video and image segmentation, extending the capabilities of its predecessor to the temporal domain. It introduces the Promptable Visual Segmentation (PVS) task, which generalizes segmentation to videos using prompts like points, boxes, or masks on any frame. SAM2 employs a streaming architecture with a memory module to track objects across frames, effectively handling appearance changes, occlusions, and motion. For single images, it functions similar to SAM, but it was trained on a newer Segment Anything Video (SA-V) dataset. Usage is identical to SAM above; the only differences are the model used (facebook/sam2.1-hiera-large) and the default and tuned hyperparameters (see Table A.5).

**Table A.5.** Default SAM2 hyperparameters and those obtained through HPO.

| Parameter | Default | Tuned |
| --- | --- | --- |
| `pred_iou_thresh` | 0.8 | 0.26 |
| `stability_score_thresh` | 0.95 | 0.70 |

#### *µ*SAM

*µ*SAM improves and extends SAM for microscopy data. It addresses the challenge of identifying objects across various microscopy modalities like light microscopy (LM) and electron microscopy (EM) by fine-tuning SAM with a new decoder, resulting in improved instance segmentation. This approach provides specialized models for LM and EM that outperform the default SAM and includes an interactive tool for annotation. *µ*SAM is the only package^9^ that needs to be installed via *conda* and is then used via

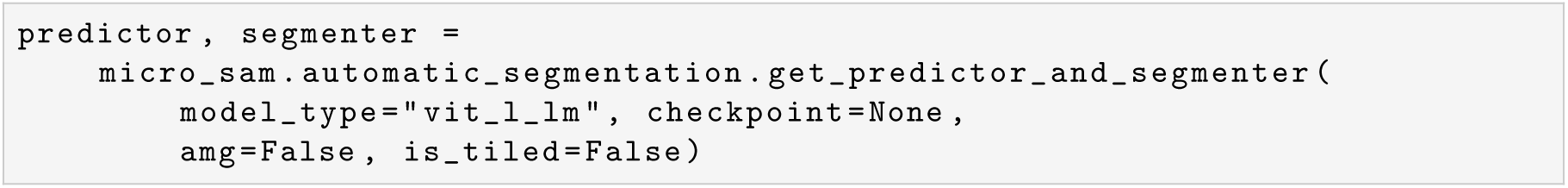

Prediction on an image are called by first computing embeddings,

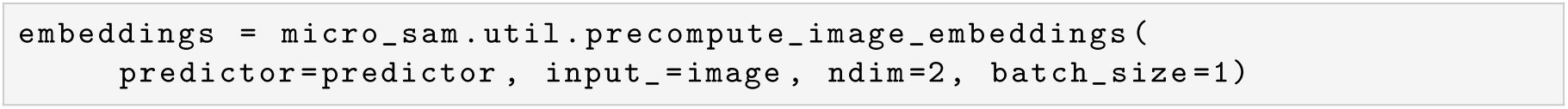

initializing the segmenter with segmenter.initialize(img, embeddings), and producing the masks using

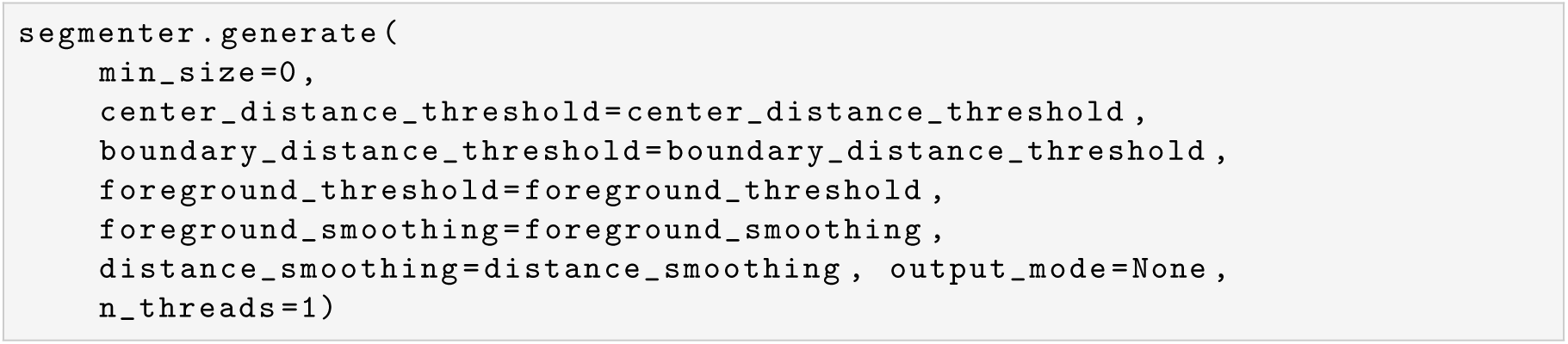

The default and tuned hyperparameters are summarized in Table A.6. Model-internal postprocessing is turned off by setting min_size=0 above (see section 4).

**Table A.6.** Default *µ*SAM hyperparameters and those obtained through HPO.

| Parameter | Default | Tuned |
| --- | --- | --- |
| `center_distance_threshold` | 0.5 | 0.89 |
| `boundary_distance_threshold` | 0.5 | 0.52 |
| `foreground_threshold` | 0.5 | 1.95 |
| `foreground_smoothing` | 1.0 | 1.11 |
| `distance_smoothing` | 1.6 | 0.73 |

#### CellSAM

CellSAM is a foundation model for cell segmentation that extends SAM to perform automated cellular instance segmentation. To overcome SAM’s limitations with dense cellular images, CellSAM employs a prompt engineering approach, using a transformer-based object detector called CellFinder to generate bounding box prompts. CellFinder and the segmentation model share the same Vision Transformer (ViT) backbone. Trained on a large, diverse corpus of cellular imaging data, CellSAM achieves state-of-the-art performance on numerous segmentation datasets. It is used via model = cellSAM.get_model(version=“1.2”)^10^ (requires setting a DeepCell access token under DEEPCELL_ACCESS_TOKEN). Prediction on an image are called via

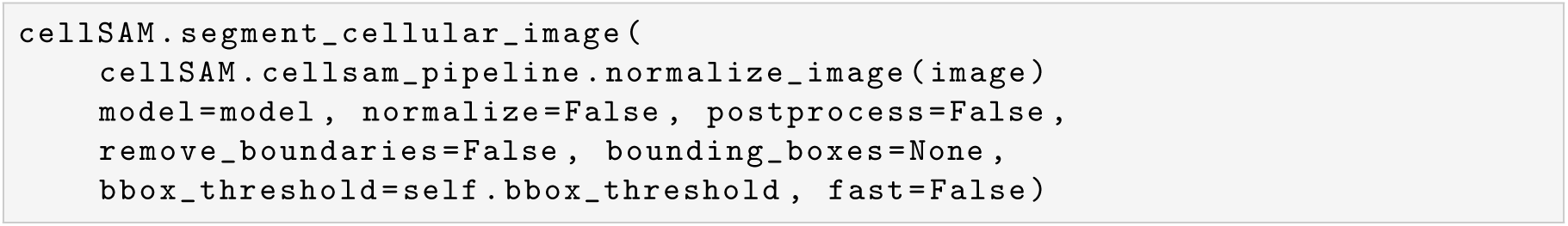

The default and tuned hyperparameters are summarized in Table A.7. We turned off all model-internal pre- and postprocessing steps by setting normalize=False, postprocess=False, remove_boundaries=False above. However, note that we do use the cellSAM.cellsam_pipeline.normalize_image as its a main part of the architecture (similar to how with SAM, we need to rescale to uint8 (see above)).

**Table A.7.** Default CellSAM hyperparameter and that obtained through HPO.

| Parameter | Default | Tuned |
| --- | --- | --- |
| cellprob_threshold | 0 | 0.37 |
| diameter | 30 | 37 |
| flow_threshold | 0.4 | 0.83 |

#### Cellpose-SAM

Cellpose-SAM integrates the pretrained transformer backbone of SAM into the Cellpose framework. This combination aims to achieve better generalization and performance, outperforming inter-human agreement and approaching a hypothetical “human-consensus” bound for segmentation quality. By leveraging the Cellpose framework’s effective loss function and post-processing with SAM’s strong inductive biases, Cellpose-SAM establishes itself as a foundation model for biological segmentation. It is used via model = cellpose.models.CellposeModel(pretrained_model=“cpsam”)^11^. Prediction on an image are called via

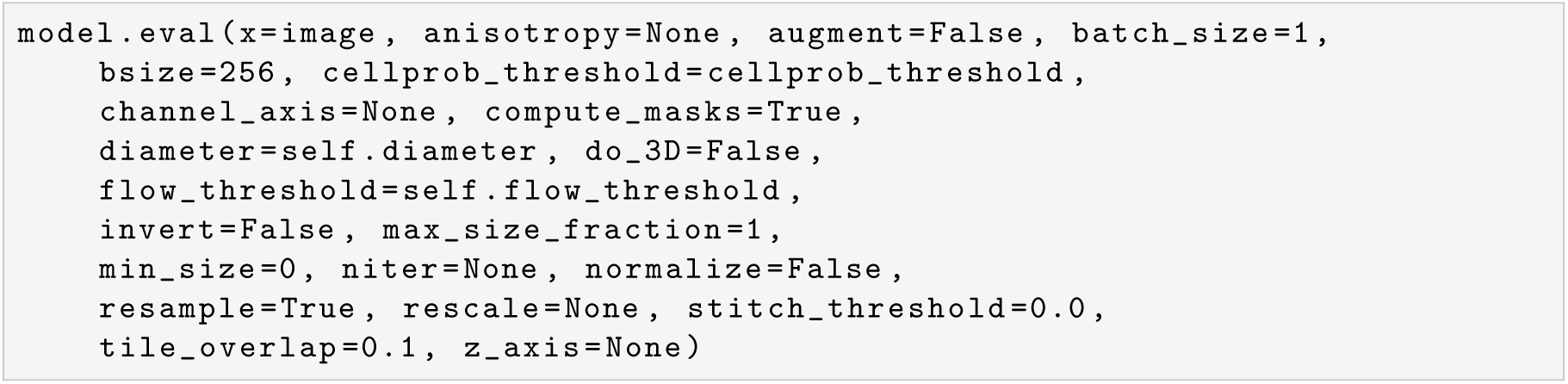

The default and tuned hyperparameters are summarized in Table A.8. We turned off all model-internal pre- and postprocessing steps by setting

**Table A.8.**
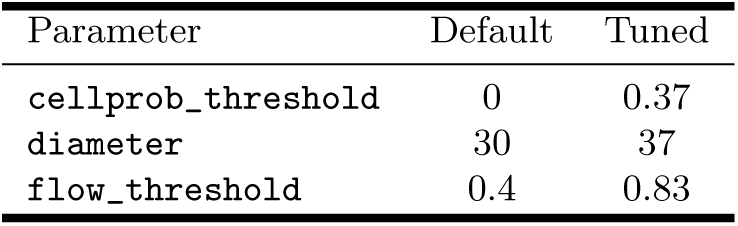
Default Cellpose-SAM hyperparameters and those obtained through HPO.

invert=False, max_size_fraction=1, min_size=0, normalize=False above.

#### SAM3

Segment Anything Model 3 (SAM3) is a unified model that detects, segments, and tracks objects in images and videos based on “concept prompts” such as short noun phrases or image exemplars. This task, termed Promptable Concept Segmentation (PCS), returns segmentation masks and unique identities for all matching object instances. The model architecture features an image-level detector and a memory-based video tracker sharing a single backbone. SAM3 is the first in the series to offer fully functional text-prompted segmentation. Identical in usage to SAM and SAM2 above; only difference is the model used (facebook/sam3) and the default and tuned hyperparameters (see Table A.9). Note that access to SAM3 on Hugging Face^12^ has to be granted.

**Table A.9.**
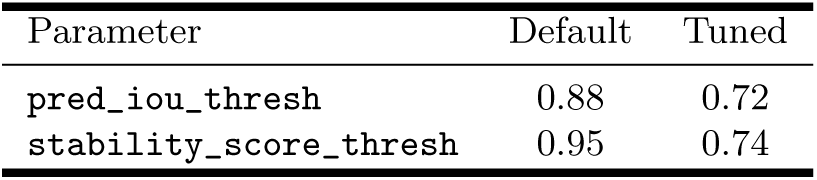
Default SAM3 hyperparameters and those obtained through HPO.

#### SAM3Text

SAM3Text uses SAM3’s text prompting capabilities instead of AIS. For that, we come up with ten different text prompts which the HPO can choose from. As a plausible default, we set thin line. The other options are elongated structure, straight black line, linear biological filament, narrow elongated line, thin red-highlighted line, thin bright-on-dark microstructure, thin structure in noisy background, small linear object, and thin linear structure among noise. The model is used via the Hugging Face transformers module,

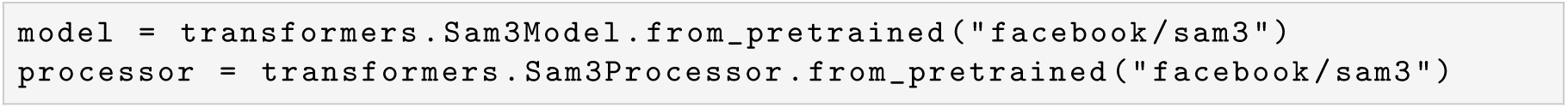

Predictions on an image are called via

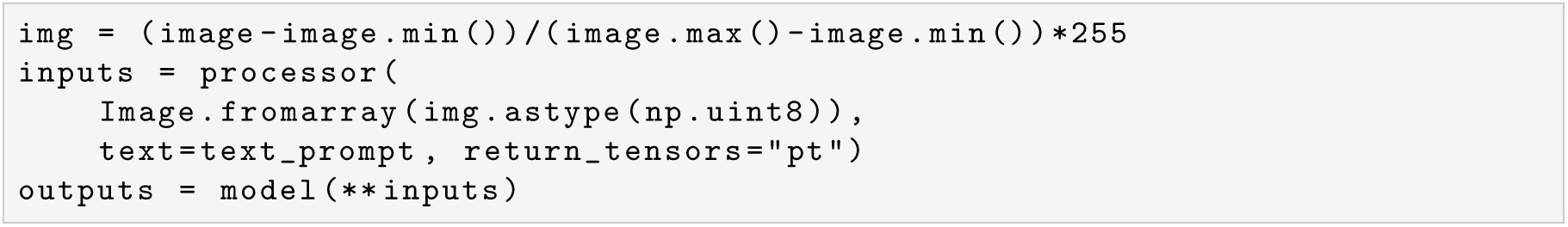

and results are given by

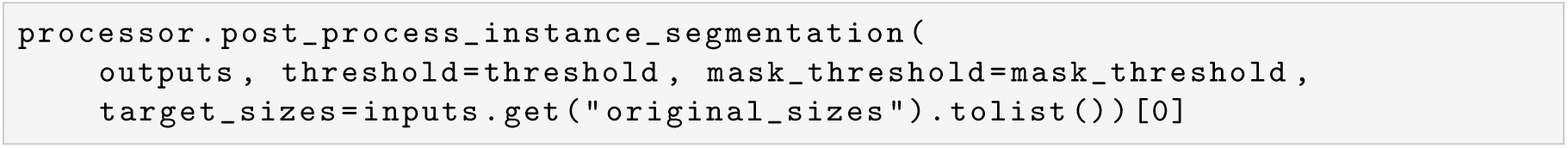

The default and tuned hyperparameters are summarized in Table A.10.

**Table A.10.** Default SAM3Text hyperparameters and those obtained through HPO.

| Parameter | Default | Tuned |
| --- | --- | --- |
| threshold | 0.5 | 0.40 |
| mask_threshold | 0.5 | 0.44 |
| text_prompt | thin line | thin line |

### A.2#Common preprocessing

All method wrappers inherit from a common base class that centralizes the preprocessing parameters grayscale, sharpen_radius, smooth_radius, percentile_min, percentile_max, clip_to_percentiles, rescale_using_percentiles, invert, and histogram_normalization. We chose the default parameters for each method such that our preprocessing pipeline reflects their internal one that we turned off (see selection of method-specific default parameters in appendix A.1). The idea behind this is that in the HPO, all methods have the same amount of preprocessing capabilities available. In Table A.11, we list both the default and tuned preprocessing parameters for each method. For more information, we refer to our implementation of preprocessing.process_image() in our codebase.

**Table A.11.** Default (Def.) and tuned preprocessing parameters obtained through HPO. Abbreviations: gryscl = grayscale, shrpn_rad = sharpen_radius, smth_rad = smooth_radius, perc_min = percentile_min, perc_max = percentile_max, clip_perc = clip_to_percentiles, rescl_perc = rescale_using_percentiles, hist_norm = histogram_normalization, f = False, t = True.

|  | `grayscale` |  | `shrpn_rad` |  | `smth_rad` |  | `perc_min` |  | `perc_max` |  | `clip_perc` |  | `rescl_perc` |  | `invert` |  | `hist_norm` |  |
| --- | --- | --- | --- | --- | --- | --- | --- | --- | --- | --- | --- | --- | --- | --- | --- | --- | --- | --- |
| Model | Def. | Tuned | Def. | Tuned | Def. | Tuned | Def. | Tuned | Def. | Tuned | Def. | Tuned | Def. | Tuned | Def. | Tuned | Def. | Tuned |
| FIESTA | `t` | `t` | 0 | 0.18 | 0 | 2.25 | 0 | 4.25 | 100 | 96.11 | `f` | `f` | `f` | `f` | `f` | `f` | `f` | `f` |
| StarDist | `t` | `t` | 0 | 8.12 | 0 | 18.10 | 0 | 0.18 | 100 | 96.71 | `f` | `f` | `f` | `t` | `f` | `t` | `f` | `t` |
| TARDIS | `t` | `t` | 0 | 7.17 | 0 | 0.36 | 0 | 2.89 | 100 | 99.99 | `f` | `f` | `f` | `f` | `f` | `f` | `f` | `f` |
| μSAM | `f` | `f` | 0 | 3.07 | 0 | 1.80 | 0 | 0.63 | 100 | 96.27 | `f` | `t` | `f` | `f` | `f` | `t` | `f` | `f` |
| CellSAM | `f` | `t` | 0 | 1.15 | 0 | 5.11 | 0 | 2.66 | 99 | 95.33 | `t` | `f` | `f` | `f` | `f` | `t` | `t` | `f` |
| Cellpose-SAM | `f` | `t` | 0 | 1.45 | 0 | 19.79 | 1 | 2.01 | 99 | 99.42 | `f` | `f` | `t` | `t` | `f` | `t` | `f` | `f` |
| SAM | `f` | `t` | 0 | 1.02 | 0 | 17.62 | 0 | 4.53 | 100 | 98.60 | `f` | `t` | `f` | `f` | `f` | `t` | `f` | `f` |
| SAM2 | `f` | `f` | 0 | 1.48 | 0 | 0.50 | 0 | 2.58 | 100 | 97.90 | `f` | `f` | `f` | `t` | `f` | `f` | `f` | `f` |
| SAM3 | `f` | `f` | 0 | 0.35 | 0 | 4.28 | 0 | 2.59 | 100 | 97.03 | `f` | `f` | `f` | `f` | `f` | `t` | `f` | `t` |
| SAMText | `f` | `t` | 0 | 0.39 | 0 | 3.41 | 0 | 3.51 | 100 | 99.46 | `f` | `t` | `f` | `t` | `f` | `f` | `f` | `f` |

### A.3#Microtubule in vitro reconstitution and microscopy

MTs were reconstituted and visualized using interference reflection microscopy (IRM) as previously described. In short, reaction chambers were constructed on PLL-PEG passivated glass slides and coverglassses functionalized with Hydroxy-*ω*-Amino polyethylene glycol (PEG) and *α*-Biotinamido-*ω*-Amino PEG [83]. The chambers were further functionalized and passivated by subsequent washes with 1% (w/v) Pluronic F-127, 1 mg/ml kappa-casein and 0.2 mg/ml Neutravidin and MTs seeds, containing fluorescently labeled, biotinylated tubulin, stabilised by GMPCPP. Dynamic polymerization reactions varied depending on experimental conditions but were carried out with 6–20 *µ*M tubulin in the presence of 1 mg/ml k-casein, 0.1%–1% (v/v) *β*-mercaptoethanol, 2.5 mM Protocatechuic Acid, 25 nM Protocatechuate-3,4-dioxygenase, 0.15%(w/v) methylcellulose in a BRB80-based buffer [84]. Functional tubulin was derived from various sources, ranging from commercially available mammalian tubulin to protozoan tubulin as described in [85]. The samples were imaged using either a Nikon Ti2 inverted widefield microscope or a Nikon Eclipse Ti-E microscope. The Ti2 system was equipped with a 50/50 beam splitter, a Nikon Plan Apochromat 60×/0.95 NA oil objective, Lumencore SpectraX LED illumination, and a pco.edge 4.2 LT HQ camera. The Ti-E system featured a 50/50 beam splitter, a Nikon Plan Apochromat 100×/1.49 NA oil immersion objective, and a Photometrics Prime 95B sCMOS camera. MTs were visualized using IRM and the seeds using a excitation with a 635 nm LED or 647 nm laser, respectively. The resulting images have a resolution of 9.0909 to 9.2308 pixels per micrometer.

### A.4#Example images from SynthMT and other available datasets

Fig. A.4 depicts examples from our SynthMT dataset (see section 4 for details about the process of creation it). Even though there is no other publicly available dataset for our task (single *in vitro* microtubules (MTs) nucleated from fixed seeds), we show samples of other datasets that are close to it. However, as noted in section 2, none of these papers validate the fidelity of their datasets and do not evaluate them on foundation models (that is, treat it as a benchmark). In Fig. A.5a, we show samples from MicSim FluoMT [19], and Fig. A.5b depicts samples from DRIFT [28].

### A.5#User interface of the study

The user interface that was used to assess the perceptual realism of SynthMT is visualized in Fig. A.6, and some sample images that appear in it are shown in Fig. A.7. We also make this code publicly available at https://github.com/DATEXIS/SynthMT-study.

### A.6#Length and curvature distributions

Besides the Kullback–Leibler divergence (KL divergence), which we report throughout the paper (e.g., in Tables 1 and 2), we visualize the actual predicted and ground-truth distributions of SynthMT for all methods in Figs. A.8 and A.9.

### A.7#Model HPO and parameter importance

For completeness, we report the progression of the Hyperparameter Optimization (HPO) (as outlined in section 4) for all evaluated methods in Fig. A.10. Recall that we use 10 random images from SynthMT and Skeleton Intersection over Union (SKIoU) as a metric to be optimized in each run. Each subplot corresponds to a distinct method and shows the best observed value for SKIoU as a function of the trial index (1000 in total). The underlying search was performed over each method’s specific (see appendix A.1) and the preprocessing hyperparameters (see appendix A.2), and their search areas are detailed in our codebase. It can be inferred that the HPO is effective for all methods, where most of them exhibit a rapid increase already within the first tens of trials.

Occasional jumps in the curves correspond to the discovery of qualitatively better hyperparameter settings, for example combinations that improve image contrast handling or regularization for faint microtubules. We use the highest validation SKIoU reached by each method as the “tuned” HPO parameters (as reported in appendices A.1 and A.2).

**Figure A.4.**
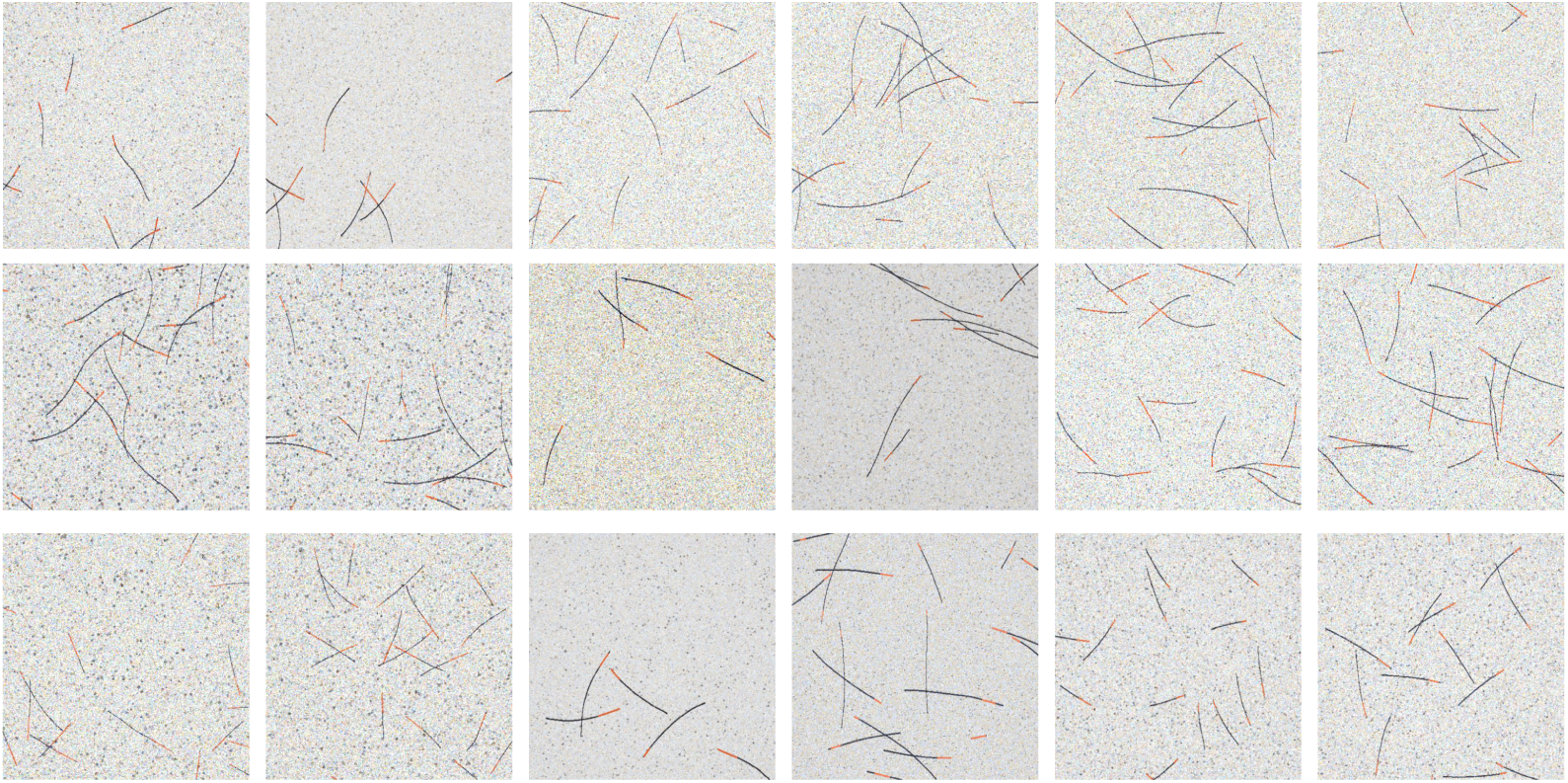
Examples from the SynthMT dataset. 18 images sampled from the synthetic SynthMT dataset. Each image depicts individual MTs growing from stabilized seeds (shown in red) under simulated interference reflection microscopy (IRM) conditions. The dataset captures natural variation in MT quantity, length, and curvature across different experimental conditions represented by 660 optimized parameter sets. The generative pipeline (Fig. 2) produces realistic imaging characteristics including background texture, vignetting effects, blur, and multiple noise sources (Poisson, Gaussian, and spatially correlated noise), and is guided by real images as depicted in Fig. 3. Every image is accompanied by pixel-accurate segmentation masks for each MTs, providing ground-truth annotations for quantitative benchmarking of segmentation methods.

The importance of each of the parameters can be inferred from the bar plots in Fig. A.11 and is calculated using f-ANOVA [86]. Except for StarDist and SAM3Text, preprocessing parameters are always most important. The parameter importance analysis further reveals that for several methods, a small subset of hyperparameters accounts for most of the performance variance. For instance, while SAM and *µ*SAM have multiple hyperparameters, their performance is predominantly influenced by a single key parameter. Similarly, the performance of CellSAM and Cellpose-SAM is largely determined by two critical parameters, although these differ between the two models. This highlights that even for methods with a larger hyperparameter space, only a few parameters are crucial for optimization in this context.

**Figure A.5.**
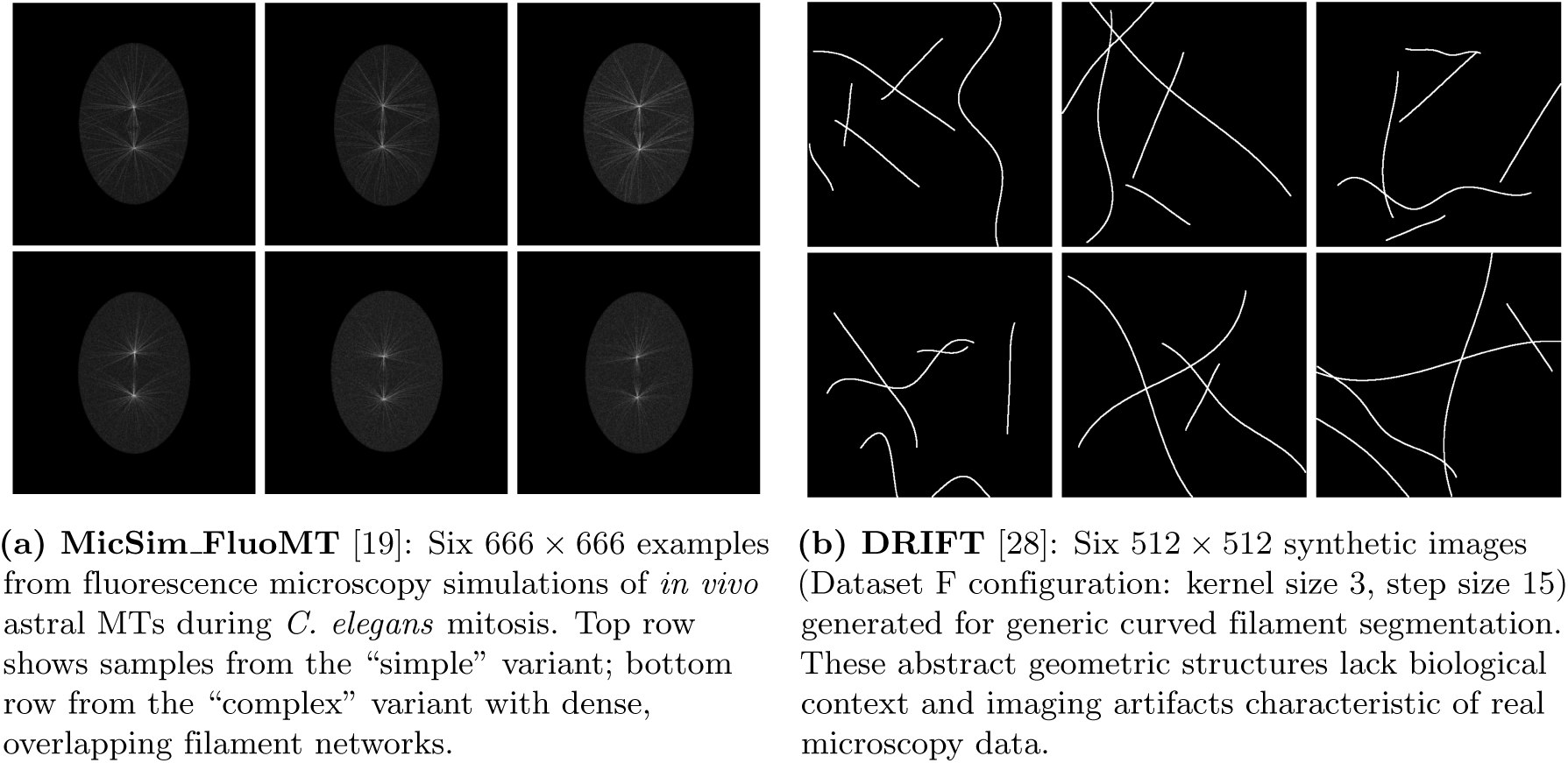
Representative samples from related synthetic MT datasets. While no publicly available dataset directly addresses our specific task of segmenting single *in vitro* MTs growing from fixed seeds under IRM microscopy, we compare against the closest available alternatives. (a) MicSim FluoMT [19] simulates *in vivo* fluorescence microscopy of astral MTs during mitosis in *C. elegans*, featuring dense, overlapping filament networks. (b) DRIFT [28] generates synthetic curved filamentous structures for general segmentation tasks. Both datasets differ fundamentally from ours in imaging modality (fluorescence vs. IRM), biological context (*in vivo* vs. *in vitro*), structural complexity (dense networks vs. individual filaments), and scope: neither provides human assessment of synthetic quality nor systematic benchmarking across foundation models, which are central contributions of SynthMT.

**Figure A.6.**
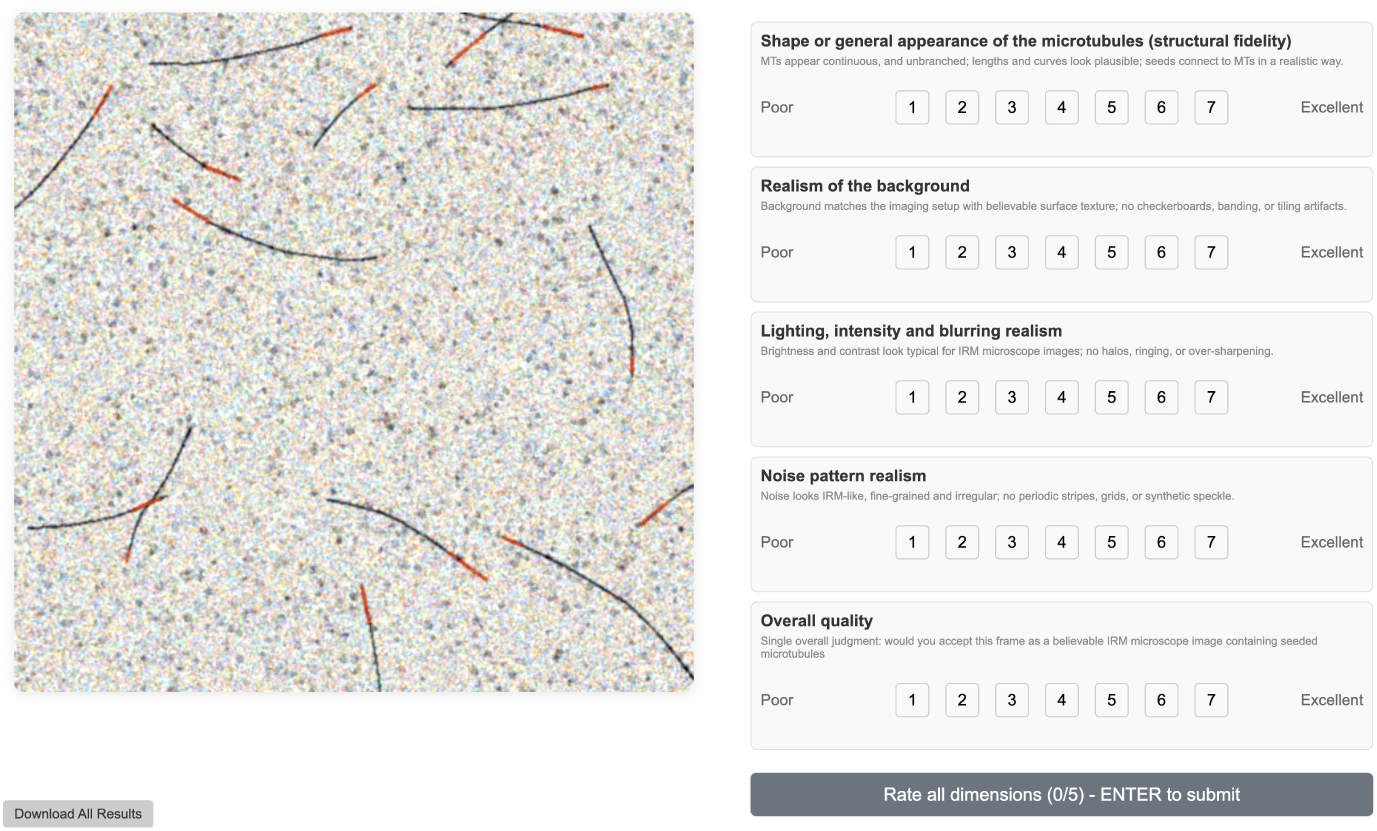
Web-based interface for expert assessment of perceptual realism of synthetic microscopy images. Participants viewed a single image at a time and rated it along five predefined dimensions using a 7-point Likert scale: structural fidelity of MTs, background realism, lighting and blurring realism, noise pattern realism, and overall quality. Images were presented in randomized order, and each image was rated exactly once per participant.

**Figure A.7.**
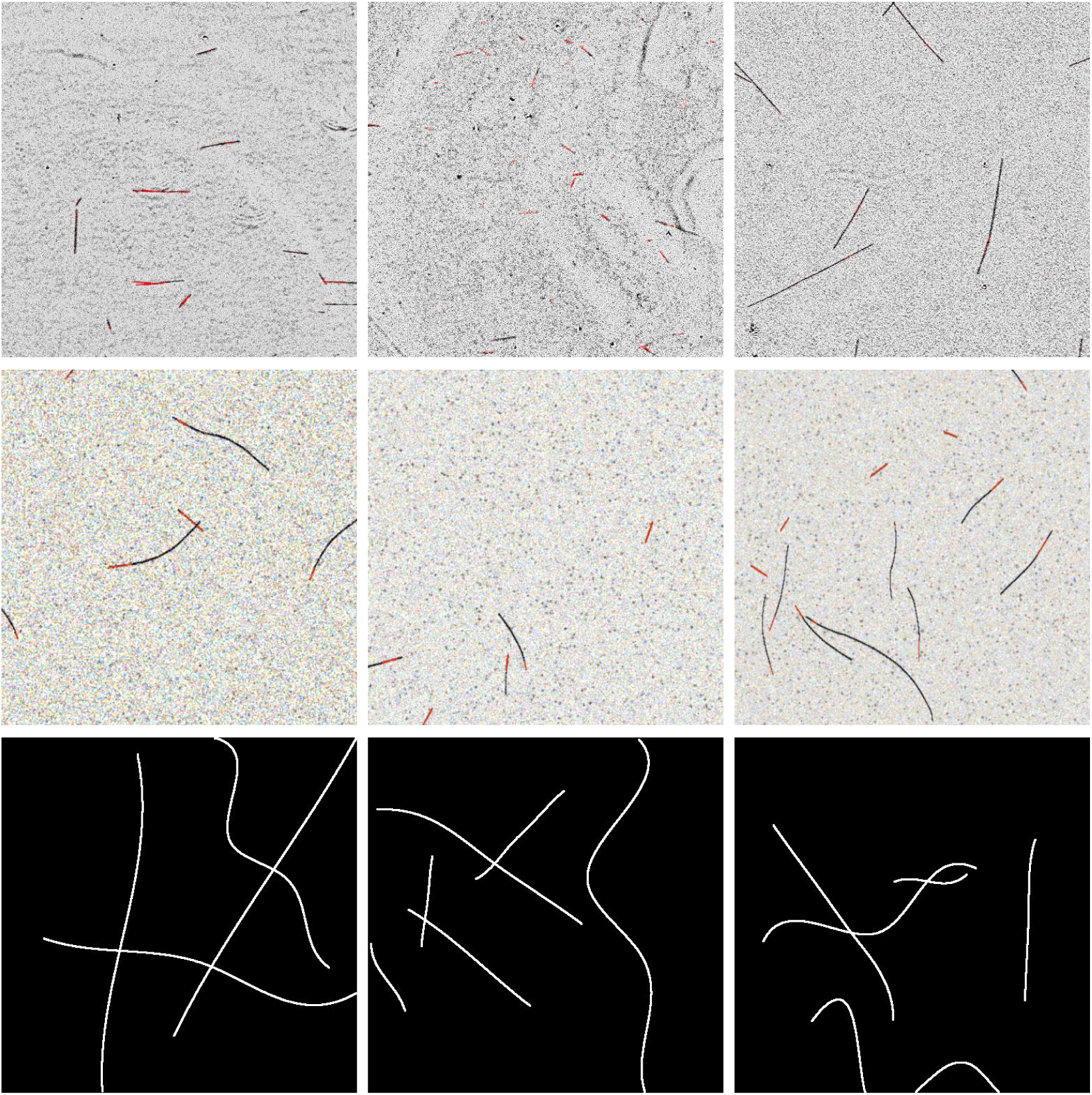
Representative images from the human validation study. Nine examples from the evaluation set shown to domain experts: three real IRM images (top row), three synthetic images from SynthMT (middle row), and three synthetic images from DRIFT [28] (bottom row). Experts rated each image on multiple quality dimensions without knowing its source, allowing us to quantitatively assess how closely SynthMT matches the appearance and characteristics of authentic microscopy data compared to existing synthetic alternatives.

**Figure A.8.**
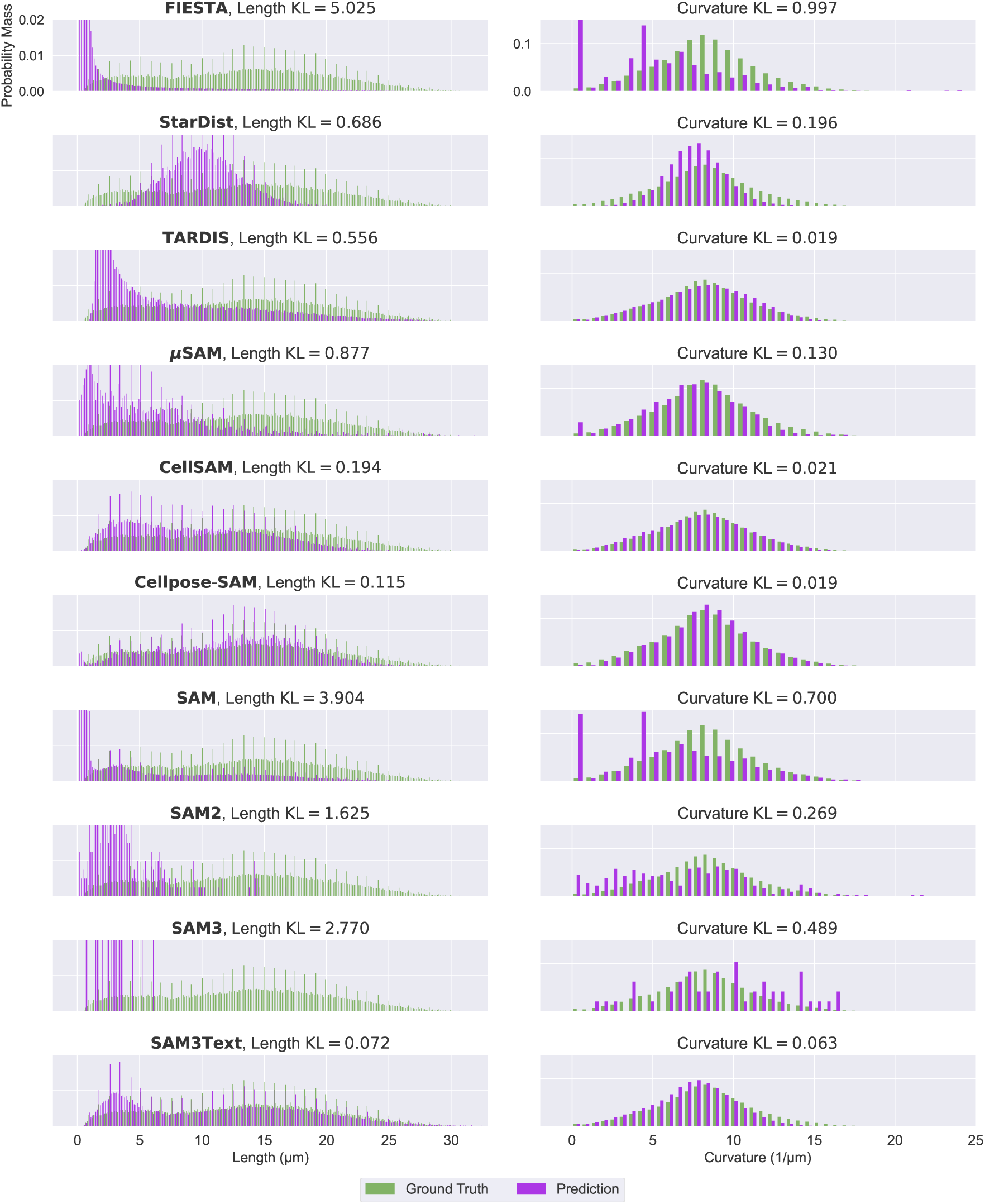
Predicted and ground-truth distributions of length and curvature for all methods with their default parameters.

**Figure A.9.**
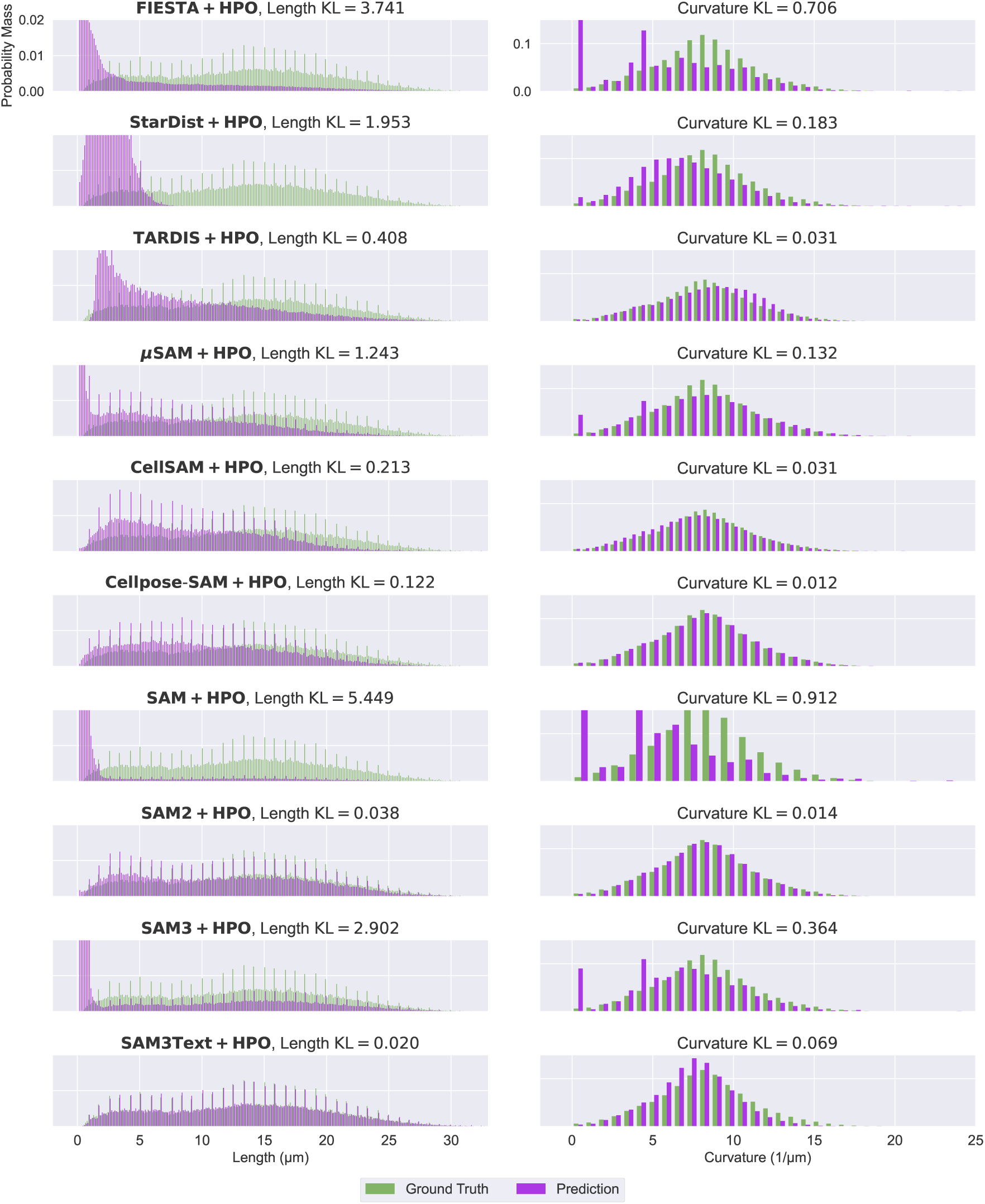
Predicted and ground-truth distributions of length and curvature for all methods with tuned parameters obtained through Hyperparameter Optimization (HPO).

**Figure A.10.**
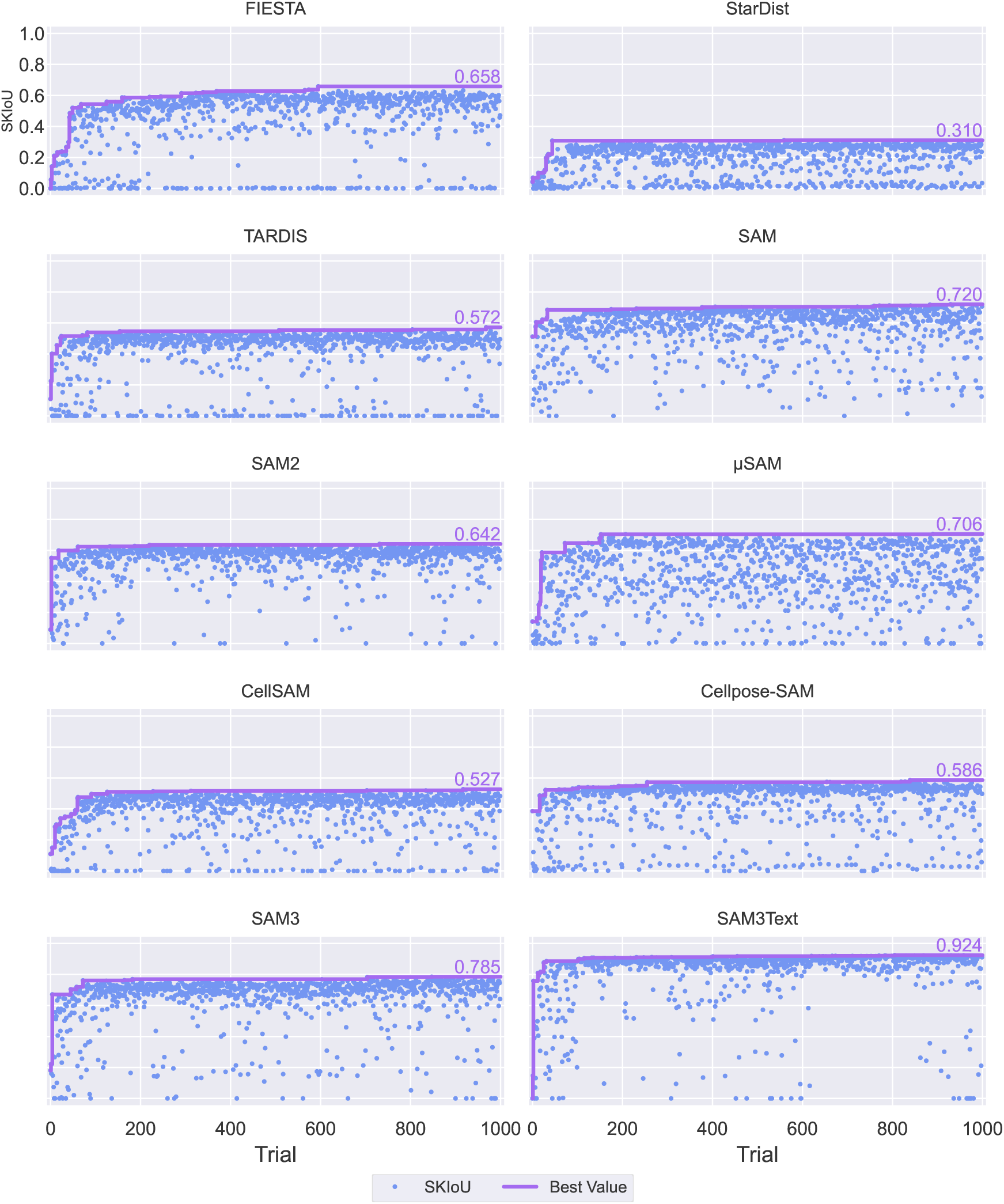
HPO trajectories highlight rapid optimization. We plot the best SKIoU value observed up to each of the 1000 HPO trials for each method, using 10 images from SynthMT for optimization. Most methods approach their optimal performance within the first 200 trials, demonstrating efficient convergence. FIESTA, which has the largest number of hyperparameters (see Table A.1), requires approximately 600 trials to converge. This illustrates both the speed of improvement and the eventual plateau in performance.

**Figure A.11.**
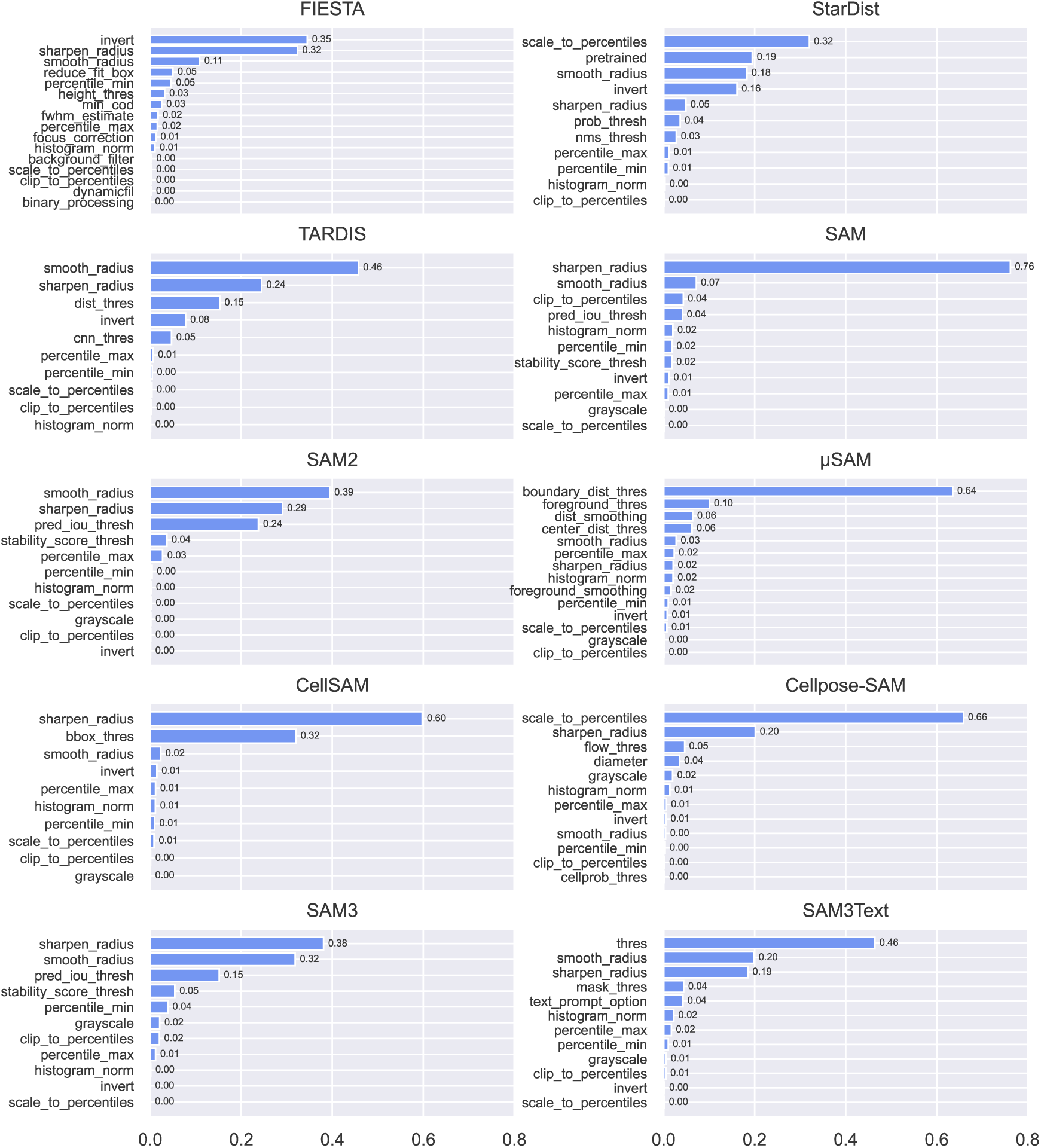
A few key hyperparameters drive method performance. Parameter importance for all models, calculated via f-ANOVA on the 10 SynthMT images used for optimization. The y-axis shows the percentage of variance explained by each hyperparameter, with higher values indicating greater impact. For most models, performance is dominated by a small number of influential parameters, which are often related to preprocessing steps.

## Footnotes

^1^As detailed in imagejdocu.list.lu/gui/process/subtract background.

^2^https://github.com/neel-dey/AnyStar/issues/4#issuecomment-3362693166

^3^See SciPy’s splprep function.

^4^https://github.com/fiesta-tud/FIESTA

^5^https://pypi.org/project/matlabengine

^6^https://pypi.org/project/stardist

^7^https://pypi.org/project/tardis-em/

^8^https://pypi.org/project/transformers

^9^https://github.com/computational-cell-analytics/micro-sam

^10^https://github.com/mario-koddenbrock/cellSAM

^11^https://pypi.org/project/cellpose

^12^https://huggingface.co/facebook/sam3

